# Tension anisotropy drives phenotypic transitions of cells via two-way cell-ECM feedback

**DOI:** 10.1101/2022.03.13.484154

**Authors:** Farid Alisafaei, Delaram Shakiba, Leanne E. Iannucci, Matthew D. Davidson, Kenneth M. Pryse, Pen-hsiu Grace Chao, Jason A. Burdick, Spencer P. Lake, Elliot L. Elson, Vivek B. Shenoy, Guy M. Genin

**Affiliations:** NSF Science and Technology Center for Engineering Mechanobiology; Department of Mechanical and Industrial Engineering, New Jersey Institute of Technology, Newark, NJ 07102, USA; Department of Mechanical Engineering & Materials Science Washington University in St. Louis, St. Louis, MO 63130, USA; Department of Biomedical Engineering, Washington University in St. Louis, St. Louis, MO 63130, USA; BioFrontiers Institute, University of Colorado Boulder, Boulder, CO 80303, USA; Department of Chemical and Biological Engineering, University of Colorado Boulder, Boulder, CO 80303, USA; Department of Biomedical Engineering, School of Engineering and School of Medicine, National Taiwan University, Taipei, Taiwan; Department of Orthopaedic Surgery, Washington University in St. Louis, St. Louis, MO 63130, USA; Washington University School of Medicine, Department of Biochemistry and Molecular Biophysics, St. Louis, MO 63130, USA; Department of Materials Science and Engineering, School of Engineering and Applied Science, University of Pennsylvania, Philadelphia, PA 19104, USA

## Abstract

Mechanical factors such as stress in the extracellular environment are known to affect phenotypic commitment of cells. However, the stress fields experienced by cells in tissues are multiaxial, and the ways that cells integrate this multiaxial information are largely unknown. Here, we report that the anisotropy of these stress fields is a critical factor triggering phenotypic transition in fibroblast cells, outweighing the previously reported role of stress amplitude. Using a combined experimental and computational approach, we discovered a self-reinforcing mechanism in which cellular protrusions interact with collagen fibers to develop tension anisotropy, which in turn stabilizes protrusions and amplifies their contractile forces. Disruption of this self-reinforcing process, either by reducing tension anisotropy or by inhibiting contractile protrusions, prevented phenotypic conversion of fibroblasts to contractile myofibroblasts.

## Introduction

Mechanical factors influence many phenotypic transitions in cells. For example, stresses, stiffnesses, and viscoelasticity affect stem cell lineage (*1–3*). In tissues, mechanical fields typically vary with direction, and cells must integrate conflicting information coming from different directions. The mechanisms for doing so are largely unknown, but are important to development, wound healing, and a number of critical pathologies.

A key example is the phenotypic transition of fibroblasts to myofibroblasts. Fibroblasts transform from a role of quiescent maintenance to one of active extracellular matrix (ECM) synthesis and contraction in response to mechanical stimuli and to mechanically associated soluble factors (*4–6*). This transformation is critical to wound healing and development, as well as to pathologies such as fibrosis (*7–11*). Controlling this transformation might enable treatment of these pathologies and advance the development of tissue-engineered systems (*12–14*). However, the fundamental mechanobiological mechanisms underlying this “activation” of fibroblasts are unclear and remain a topic of debate (*15, 16*).

This fibroblast to myofibroblast transformation is evident in reconstituted tissues that serve as simple three-dimensional (3D) culture systems. Fibroblasts are round, small, and inactive over the first hours after being embedded within a 3D collagen matrix (early stage of cell-matrix interaction, fig. S1A), but become polarized, spread, and contractile thereafter (late stage of cell-matrix interaction, fig. S1B). Although these later stages, when cells have adapted to and remodeled their mechanical microenvironment, have been well studied (*17, 18*), the mechanisms through which cells transform to these stages have yet to be elucidated.

In these late stages, cells sense and respond to mechanical stimuli and perturbations in their mechanical microenvironment (*19–23*). Upregulation of actomyosin-based contractile forces and increased cytoskeletal stiffness can result from changes to the ECM (*24–26*), including changes made by the cells themselves as they pull, align, and stiffen nearby ECM (*27–32*). Late-stage cell-ECM interactions affect migration, differentiation, proliferation, and signaling (*33–35*), and perturbation of these interactions can disrupt physiological processes (*36, 37*). However, much less is known about how the cell’s early-stage environment transforms the cells and guides its activation, or about how the cell transforms its early-stage environment. We therefore studied early-stage interactions between human dermal fibroblasts and their 3D fibrous matrix to quantify and model this two-way feedback loop, and learn to control fibroblast transformation. Our results revealed that the directionality of mechanical signals from the ECM at the early-stage plays a more important role than the intensity of these signals in the activation of fibroblasts.

### Microtubule-driven cell protrusions guide ECM alignment

We began by chronicling the first stages of ECM remodeling, a seemingly one-way feedback process in which cells guide ECM alignment. Isolated fibroblasts cultured within randomly oriented type I collagen were initially small and round (Fig. 1A, left panel), but over several hours developed patterns of protrusions that aligned collagen surrounding the protrusions (Fig. 1A, right panel), with more collagen accumulation around the bases of the protrusions, and less on other regions of the cell body (Figs. 1B and S2). Each protrusion was driven by a central microtubule (fig. S3A and S4) (*38*). Preventing growth of protrusions, by pre-treating fibroblasts with the microtubule inhibitor nocodazole before embedding and culturing them in a collagen-nocodazole mixture, prevented cells from developing polarity (fig. S3B) and reduced the degree and spatial extent of ECM alignment (Fig. 1C). Conversely, applying nocodazole 12-24 hours after embedding fibroblasts into the ECM (post-treatment) increased collagen remodeling and alignment as protrusions retracted (Fig. 1D), consistent with our previous reports of microtubule disruption increasing cell contractility in 2D culture through phosphorylation of myosin light chain (*39*). These observations highlight a central role for protrusions in driving ECM alignment (Fig. 1E).

**Fig. 1.**
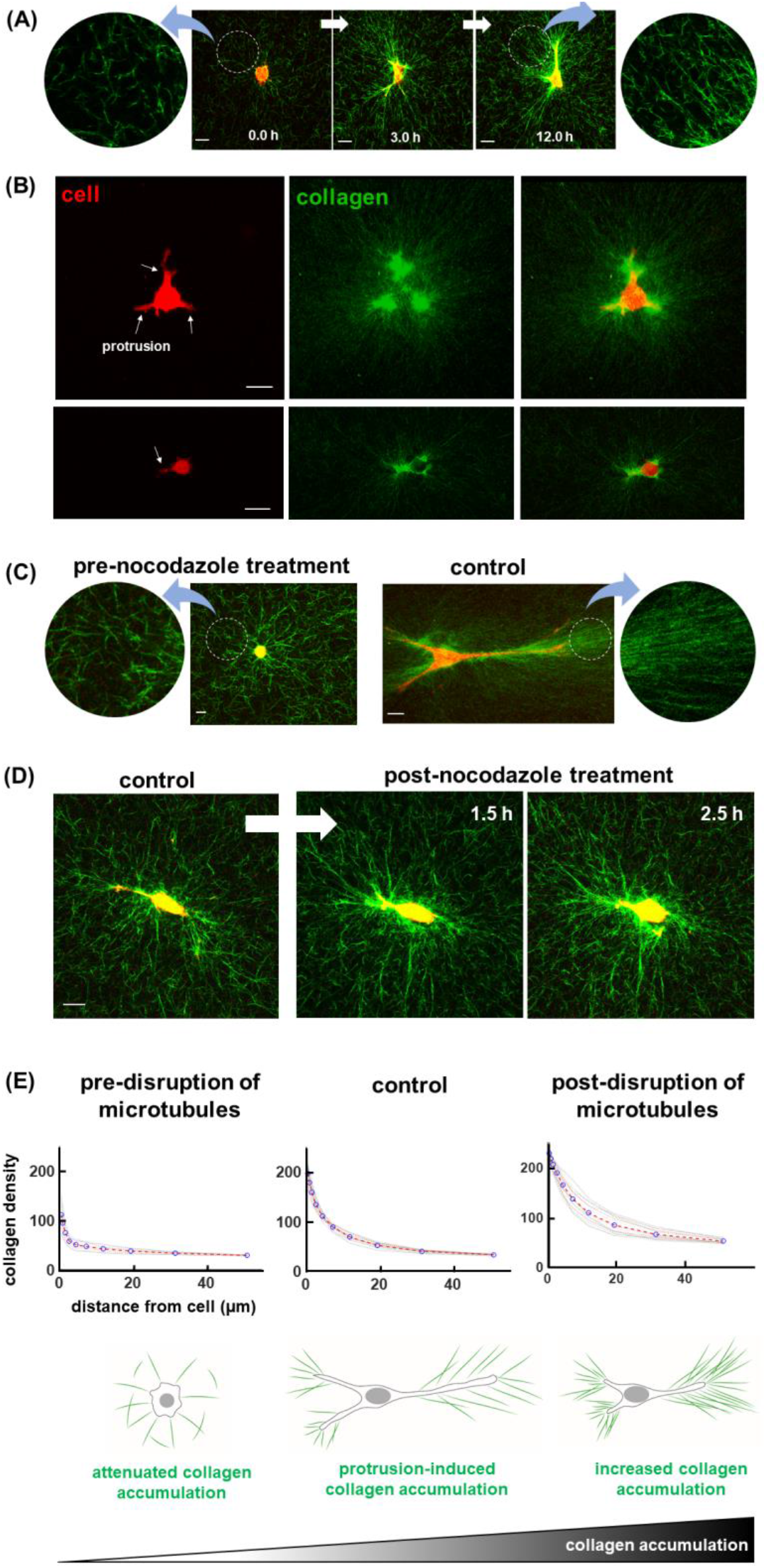
Fibroblasts align collagen and generate an anisotropic mechanical environment via microtubule-driven protrusions. **(A)** Fibroblasts were initially small and round within a 3D collagen network with randomly distributed fibers (isotropic mechanical field). Within a few hours, fibroblasts formed long protrusions and align collagen fibers along the protrusion direction. **(B)** Independent of the cellular shape and size, collagen was mainly aligned by cell protrusions, and not the cell body, indicating the importance of cell protrusions in the alignment of collagen in the early stages of cell-matrix interaction. **(C)** Inhibition of cell protrusions through pre-treatment with nocodazole (disruption of microtubules before embedding the cell within the matrix) significantly reduced collagen alignment. **(D-E)** Opposite to the results from pre-disruption of microtubules, post-disruption of microtubules caused retraction of protrusions and higher collagen alignment. Scale bar: 20 µm

### ECM alignment guides cell protrusions

ECM anisotropy (*40*) similarly provided apparently one-way guidance to cell protrusions (Fig. 2, original data from (*41*)) (*42–45*). When cultured either atop or within collagen matrices with pre-aligned fibers (see Figs. S5 and S6 for more details), fibroblasts predominantly adopted the polarity of the ECM, independent of matrix dimensionality (2D or 3D) or matrix thickness (Figs. 2A-2D). Similar to collagen alignment and accumulation (fig. S7), cell alignment also required actomyosin contractility, as evident from disruption of actomyosin contractility using the rho inhibiter Y-27632 (Fig. 2E). Thus, fibroblasts required actomyosin-driven cytoskeletal tension to become polarized, and the anisotropic mechanical environment induced by matrix fiber alignment promoted this polarization.

**Fig. 2.**
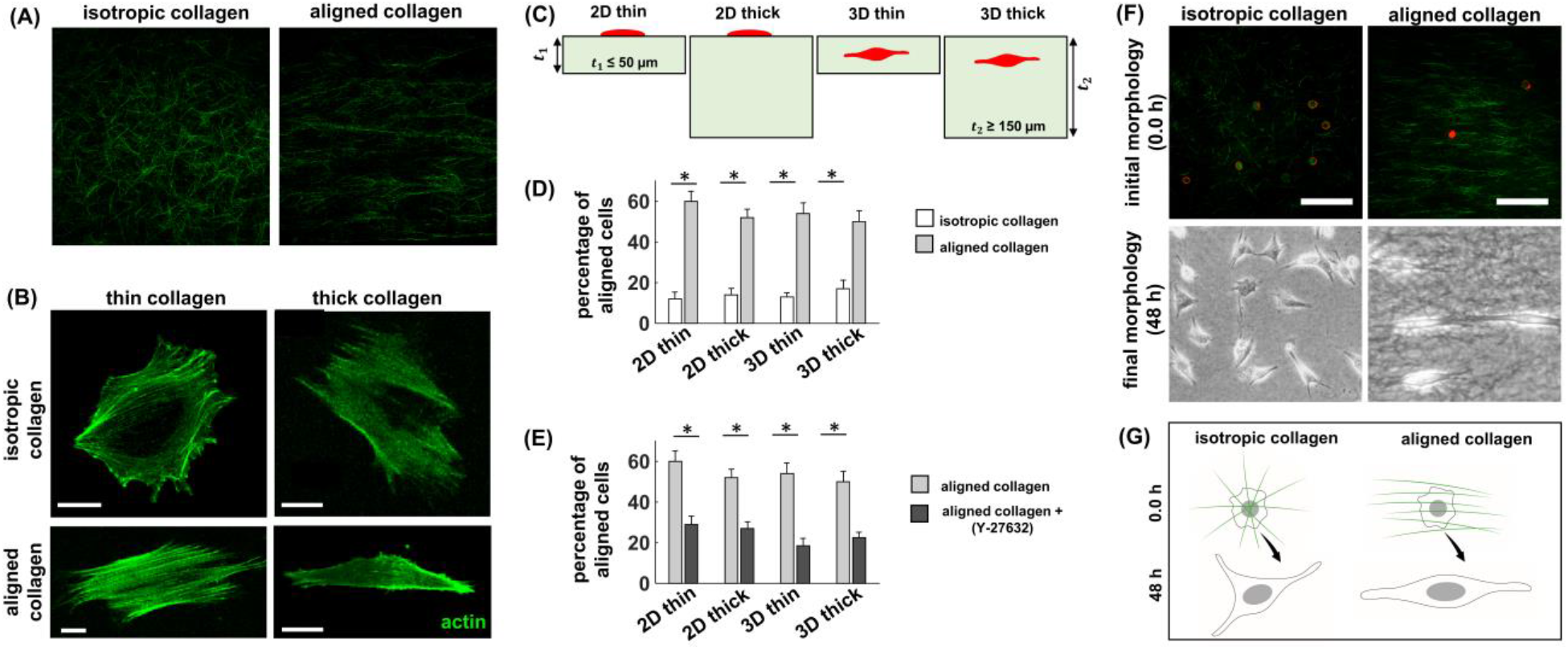
An anisotropic mechanical environment generated by collagen fiber alignment promotes the growth of protrusions in the direction of fiber alignment and prevents protrusions from growing in any other direction. **(A-D)** Using a microfluidic device, and without applying any external forces, we generated aligned and isotropic fibers to form 2D and 3D collagen matrices with both thin and thick matrix thicknesses. Fibroblasts became aligned and polarized along the direction of collagen alignment independent of matrix thickness or matrix dimensionality. **(E)** This restriction of spreading area to the direction of collagen alignment is governed by the actomyosin contractility of cells where reduction of contractility (upon Y-27632 treatment) significantly decreased the percentage of aligned cells. **(F-G)** Fibroblasts were initially round and small in both isotropic and aligned matrices, but became spread and polarized over time in both groups. The process of spreading of fibroblasts was initiated by the formation of protrusions in both groups, however, the cellular protrusions in aligned matrices were longer and were formed only in the direction of collagen alignment. Error bars represent the standard error of proportions with *n* = 106-246 in (D) and *n* = 82-184 in (E). Scale bar: 20 µm

To identify the mechanisms by which anisotropy of the mechanical environment promoted polarization, we followed isolated fibroblasts within both isotropic and aligned matrices over time. Fibroblasts in both groups were initially round and small (Fig. 2F), then tended to polarize over time, with polarization accompanied by formation of protrusions and development of cytoskeletal tension that deformed the surrounding matrix. Cellular protrusions in aligned matrices were longer and formed only in the direction of collagen alignment (Fig. 2F). This motivated a hypothesis that local mechanical anisotropy promoted cellular protrusion growth in the direction of collagen alignment, while inhibiting protrusions in other directions (Fig. 2G).

### ECM mechanical anisotropy alone is sufficient to trigger anisotropy of cell protrusions independent of ECM chemical properties

To test our hypothesis, we developed electrospun, cross-linked, synthetic fibrous hydrogels with fibers either aligned or distributed isotropically. These hydrogels are composed of chemically modified hyaluronic acid and adorned with adhesive ligands containing peptides (RGD) to facilitate cell attachment and spreading. We have previously shown that hydrogel fibers are stable over time and mimic the mechanics and microstructure of natural fibrous matrices (*46–48*) with minimal degradation even in the presence of matrix metalloproteinases (*49*). These properties of synthetic fibrous hydrogels enabled testing of our hypothesis while controlling against possible roles of chemical enzymatic changes associated with cross-linking or degradation of natural collagen-based matrices. Similar to their responses within collagen matrices (Figs. 2F-2G), initially round fibroblasts polarized after 24 hours in both isotropic and aligned hydrogels (Figs. 3C, 3D, and S8). As with collagen matrices (Figs. 2F-2G), polarization was driven by cell protrusions in both groups (Figs. 3A-3B). However, while cell protrusions in isotropic hydrogels were distributed randomly after 2 hours (Fig. 3F), cell protrusions in aligned hydrogels aligned with matrix fibers even at the early stage of cell-ECM interaction (Fig. 3E). Results, therefore, supported our hypothesis that local mechanical anisotropy promotes cellular protrusion growth in the direction of collagen alignment, while inhibiting protrusions in other directions (Movie 1).

**Fig. 3.**
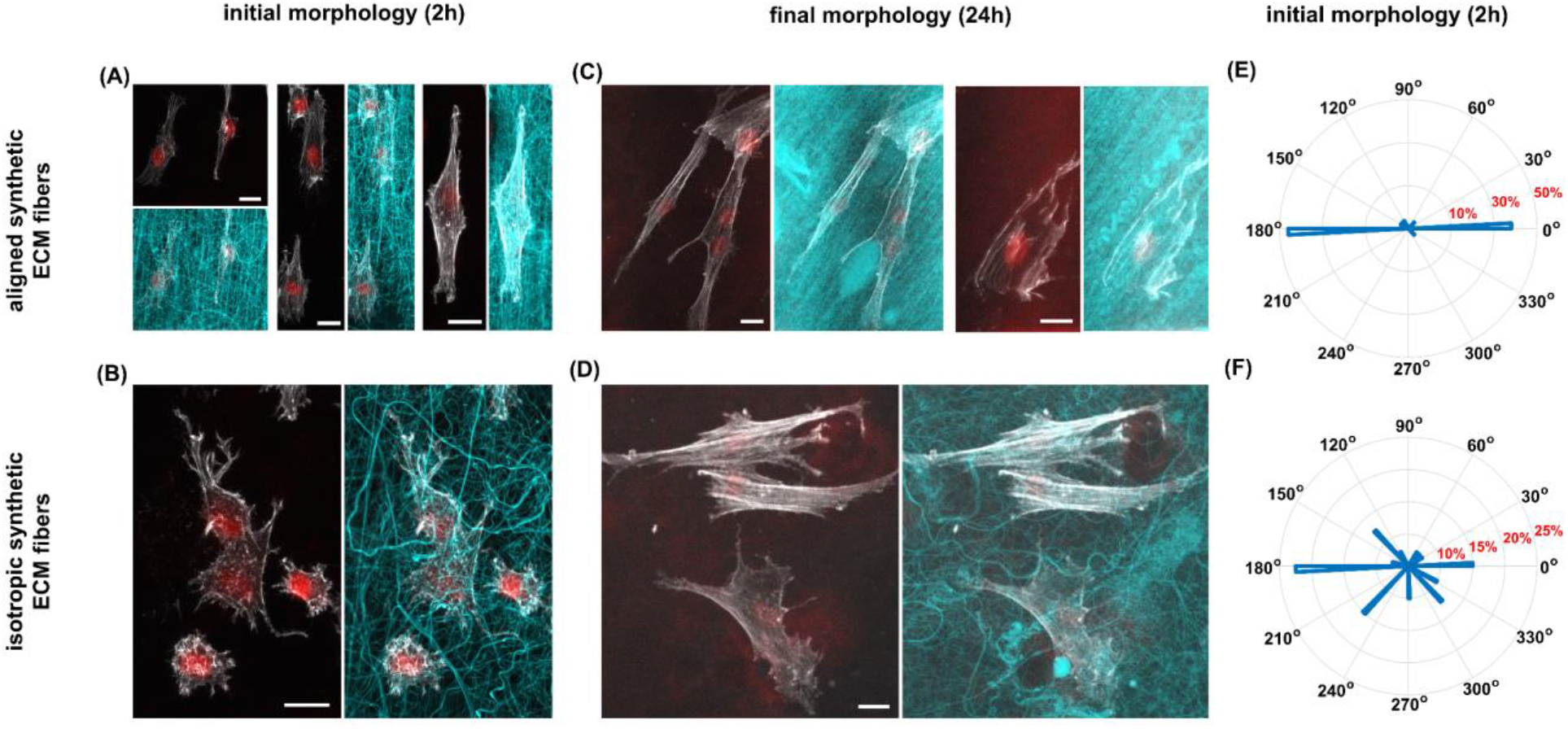
Culturing fibroblasts within 3D synthetic hydrogels confirms that an anisotropic mechanical environment alone is sufficient to trigger anisotropy of cell protrusions independent of matrix fiber chemical properties. We tracked fibroblast morphologies over time in both aligned **(A** and **C)** and isotropic **(B** and **D)** synthetic fibrous matrices which, unlike collagen matrices, are not degradable even in the presence of collagenase. Unlike cells in isotropic matrices, cell protrusions in aligned hydrogels were only formed in the direction of matrix fiber alignment from the beginning of cell interaction with the matrix **(E** and **F)**. Scale bar: 20 µm

### Two-way feedback exists between cell protrusions and ECM alignment

The two forms of one-way feedback described above occurred in two opposite directions: fibroblast protrusions determined the local alignment of matrix fibers (Fig. 1), and pre-aligned matrix fibers promoted alignment of cellular protrusions (Figs. 2 and 3). We hypothesized that these one-way factors are intertwined through a two-way feedback loop.

To test this hypothesis, we first investigated how protrusions and ECM alignment evolved after fibroblasts were embedded within collagen matrices. Initially, round fibroblasts (Fig. 4A(i)) extended small protrusions after about an hour (Fig. 4A(ii)). As shown in a single confocal slice with three small protrusions (Fig. 4A(ii)), initial perturbations in collagen alignment around retracting protrusions were amplified as the protrusions subsequently extended along the direction of collagen alignment (Fig. 4A(iii)) to lengths greater than observed in Fig. 4A(ii). Retraction of protrusions followed by extension to still greater lengths continued (Fig. 4A(iv)), with collagen alignment increasing over time. Comparison of the final collapsed z-stack (Fig. 4A(v)) to the initial (Fig. 4A(i)) showed extensive ECM remodeling. Results supported the hypothesis of two-way feedback: cell protrusions align collagen fibers in their vicinity, and these aligned fibers stabilize directions of protrusions (Fig. 4B).

**Fig. 4.**
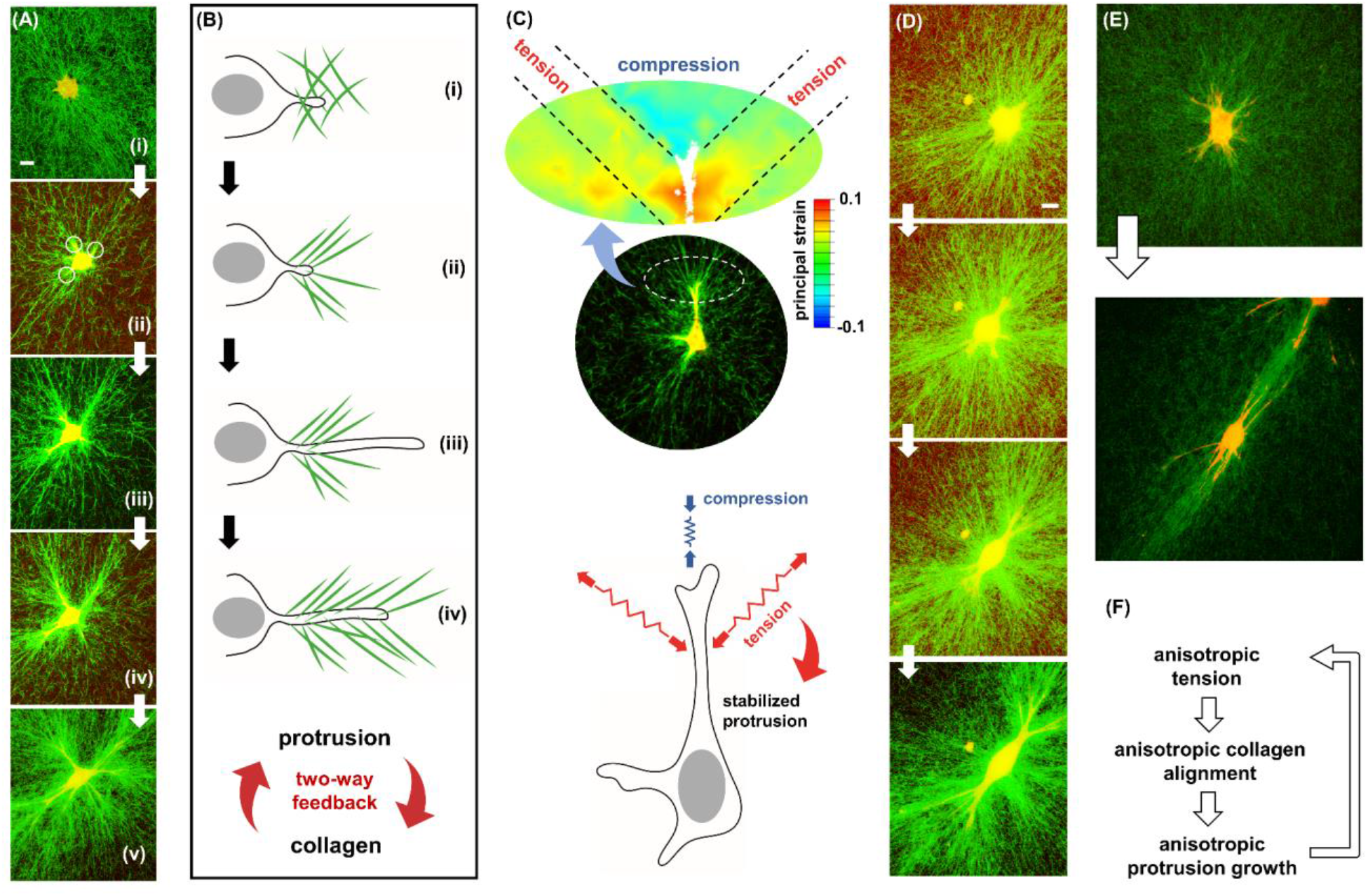
Two-way feedback between collagen alignment and protrusion stabilization causes fibroblasts to spread, polarize, and contract. **(A-B)** Cell protrusions align collagen fibers in their vicinity, and the aligned fibers in turn stabilize the protrusions leading to the growth of the protrusion in the direction of collagen alignment. **(C)** Protrusion-induced collagen alignment generated resistive tensile stresses at the cell-matrix interface, which in turn stabilized the protrusion by preventing it from being fully retracted. The stabilized protrusion grew from the tip (compressed region), leading to a periodic extension and retraction of protrusions and subsequently collagen accumulation. **(D-F)** The two-way feedback between protrusion stabilization and collagen alignment enables fibroblasts to polarize in the direction of collagen alignment and generate anisotropic stress fields.

This two-way feedback gradually amplified as effects accumulated over cycles of extension and retraction. As retraction and extension of protrusions are associated with pulling and pushing the surrounding collagen fibers, respectively, we next asked how the cyclic extension and retraction of protrusions can lead to accumulation and alignment of collagen. Note that retraction and extension of protrusions can have opposite effects on the alignment and accumulation of collagen fibers. As protrusions retract, they pull on and align collagen fibers, while the re-extension of the protrusions can fully dissipate the collagen alignment unless one of the two following scenarios (or combination of them) occurs: (i) collagen fibers undergo plastic deformation during protrusion retraction, (ii) protrusions re-extend without releasing the aligned collagen fibers. In the first scenario, the protrusion releases the tension on the aligned collagen fibers to re-extend, however, collagen fibers remain aligned and accumulated as they have undergone plastic (permanent) deformation during the protrusion retraction. In the second scenario, the protrusion re-extends while it maintains the tension in the aligned collagen fibers and keeps them aligned. In what follows, we examine both scenarios.

To test the degree to which plastic deformation (*50–52*) of the ECM contributed, the actomyosin contractility inhibitor cytochalasin D was added while individual cells were tracked. Collagen alignment and accumulation relaxed to a degree when actomyosin contractility was inhibited, but not completely, indicating both elastic and plastic deformation of the ECM (fig. S9). Application of the actomyosin contractility agonist calyculin A, which caused cells to tear away from the ECM, resulted in similar responses (fig. S10), again suggesting both elastic and plastic deformations of the ECM. Thus, effects of fibroblasts on ECM alignment and compaction were partially sustained through cycles of process extension and retraction by plastic deformation in the matrix.

To test whether protrusions can re-extend while maintaining the tension in the aligned collagen fibers (second scenario), we used a 3D strain mapping method to visualize the strain field within the ECM during periodic retraction and extension of protrusions, as has been performed previously for this class of tissue construct (*38, 53, 54*). As we have reported before (*38*), protrusions pulled on ECM from their lateral flanks and compressed it in a cone ahead of the protrusion tip (Figs. 4C and S11), indicating maintenance of lateral tension during protrusion growth. Thus, cytoskeletal tension, critical to cell alignment and polarization (cf. Fig. 2E), was maintained throughout the cyclical processes of extension and retraction. Taken together, our 3D traction force microscopy results revealed the key role of tensile forces in the two-way feedback mechanism; the protrusion-induced collagen alignment generated local tensile forces in the ECM which in turn stabilized the protrusion by preventing it from being fully retracted (Fig. 4C). In return, the stabilized protrusion grew from the tip leading to another cycle of extension and retraction (Figs. 4B-4C).

### The two-way feedback mechanism generates anisotropy of the stress field and cell polarization

We next asked what regulated the development of polarity during this positive feedback loop between protrusions and collagen fibers. The positive feedback led to a periodic extension and retraction of protrusions, culminating in a previously observed plateau in collagen accumulation over several hours (fig. S12) (*38*). Concomitantly, fibroblasts transformed from an initially round, small, and inactive state with protrusions in random directions to a spread, polarized, and active state with protrusions in a preferential direction (Figs. 4D-4E). As the latter state is associated with the generation of an anisotropic stress field (Fig. 4F), we asked whether stress anisotropy is essential for the transformation and activation of fibroblasts. To test the hypothesis, we integrated a theoretical model (*55*) with an experimental system in which stress anisotropy could be spatially controlled within the same tissue.

### Disruption of anisotropic stress fields prevents fibroblast activation

The theoretical model (see (*56*) and Supplementary Information) accounted for how cells increase their contractile forces in response to tension through a feedback loop between the density of phosphorylated myosin motors (*ρ*_*kk*_) and the tension generated in the cytoskeleton (*σ*_*kk*_) (Fig. 5A). The model thus predicted that cell contractility (denoted by the average of *ρ*_*kk*_) increases with external resistance to contraction arising from increases in either cell-substrate contact area (fig. S13A) or substrate stiffness (fig. S13B): these factors elevated cytoskeletal tension, which in turn led to more phosphorylated myosin motors and generation of higher contractile forces (*55, 57, 58*).

**Fig. 5.**
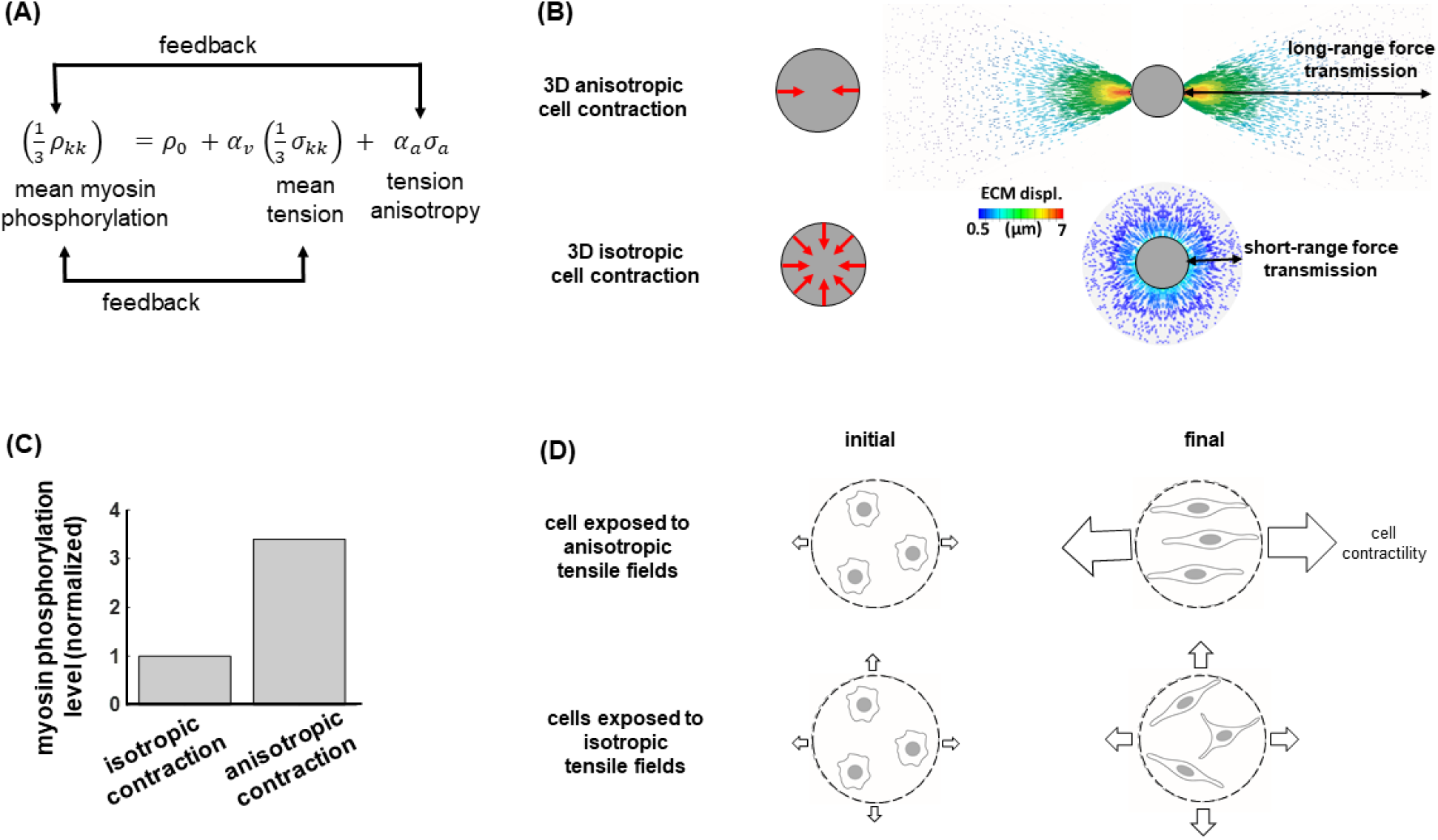
Mathematical model of how the magnitude and anisotropy of stress regulate actomyosin contractility. **(A)** Cell contractility (denoted by the average density of phosphorylated myosin motors) increases with increasing external resistance from the matrix which can be caused by increasing cell-matrix contact area or increasing matrix stiffness. However, in addition to the external resistance from the matrix, the polarity of the tensile field (tension anisotropy) generated in the matrix also impacts the level of cell contractility. **(B-C)** Cells with polarized contraction exhibit higher actomyosin contractility and generate higher contractile forces. **(D)** Similarly, the model predicts that cells exposed to anisotropic stress fields, which can be caused by the tissue boundary conditions, exhibit higher actomyosin contractility and generate higher forces compared with cells exposed to isotropic stress fields. The model prediction is experimentally validated in Fig. 6.

To evaluate the effects of the polarity of the tensile field (tension anisotropy) on cell contractility (*55*), an additional model term was considered (Fig. 5A). This enabled prediction of experimental observations that cells exhibit lower actomyosin contractility when subjected to isotropic tension (equal in all directions), as arises when cells are confined to circular regions, than they do when subjected to anisotropic tension, as arises when cells are confined to regions that enable them to adopt polarity (fig. S14) (*55*).

We next extended the theoretical model to study how, in 3D, stress anisotropy influences contractility and can trigger activation of fibroblasts. To this end, we simulated polarized and isotropic contraction of a cell within a 3D collagen fibrous network (Fig. 5B), the latter modeled by a constitutive law that was previously validated using 3D particle tracking microscopy for various type I collagen matrices, including the 1.0 mg/mL ECM used in this study (*32*). Coupling of the matrix and cell models enabled predictions of how fibroblasts interact with ECM in both isotropic and anisotropic stress fields. In the isotropic case, cells contracted with the same initial contractility *ρ*_0_ in all directions, while cells in the polarized case contracted with the initial contractility *ρ*_0_ only in the direction of polarity. Because the effective cell-matrix contact area is significantly higher for isotropic contraction due to the focused regions of cell-matrix adhesion in the polarized case (fig. S15), isotropic contraction is expected to generate higher cytoskeletal tension and thus greater phosphorylation of myosin motors. However, the model predicts that the effect of tension anisotropy in polarized contraction overpowers the effect of higher effective contact area in isotropic contraction. Polarized contraction generates an anisotropic stress field which is predicted to significantly promote cellular actomyosin contractility (higher *ρ*_*kk*_) and cell-generated contractile forces (Figs. 5C-5D), which in turn cause higher collagen alignment in the direction of the contraction (horizontal direction in Fig. 5B), and long-range transmission of these forces through the ECM.

To test our hypothesis that tension anisotropy promotes actomyosin contractility, we developed an integrated experimental system and theoretical model of a cruciform specimen in which the initial fibrous structure was spatially uniform, but the stress field within a single specimen varied spatially from isotropic to anisotropic (Fig. 6A) due to geometrical constraints at the tissue boundaries (*2, 59–61*). Tissue contraction led to uniaxial (anisotropic) tension in the arms and equibiaxial (planar isotropic) tension in the center and close to the arm ends (Fig. 6A). Our combined cell-ECM theoretical model predicted that fibroblasts in the arms, sufficiently far from the tissue center and the arm ends, experienced anisotropic tension along the direction of the arm and had lower effective cell-matrix contact areas compared with the cells at the tissue center, and that they exhibited higher actomyosin contractility (*ρ*_*kk*_). Predictions were in excellent agreement with F-actin staining of the cruciform specimens after three days of culture, a proxy for the actomyosin contractility level (Fig. 6B). Predictions and experiments both showed that the level of contractility increased with distance from the tissue center until it reached its maximum, and then decreased towards the ends of the arms (Fig. 6B) as expected due to the loss of tension anisotropy near the constrained ends (Fig. 6A).

**Fig. 6.**
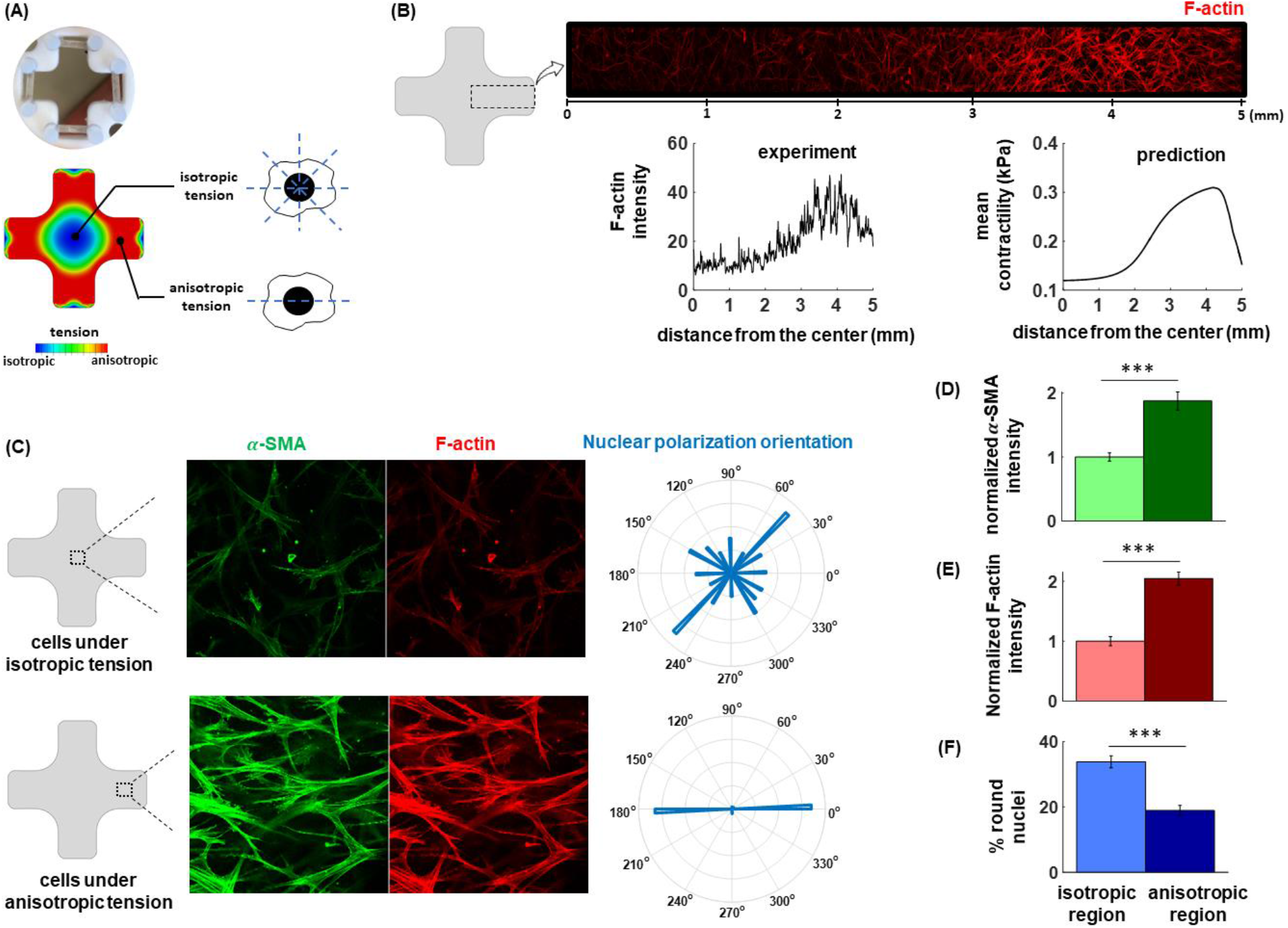
Promotion of actomyosin contractility is driven by stress anisotropy, not stress. **(A)** Cell-induced contraction of cruciform tissue specimens caused cells in center of specimens to experience equibiaxial, isotropic tension (the same in all in-plane directions), and cells in the arms to experience anisotropic (polarized) tension. **(B)** Isotropic stress fields at the center and around the arm ends prevented promotion of actomyosin contractility compared to anisotropic fields of the same peak principal stress, as shown by lower F-actin levels. **(C-F)** Cells experiencing anisotropic stress fields aligned in the direction of tension anisotropy and had more elongated nuclei than cells exposed to the isotropic stress fields. Cells subjected to anisotropic stress fields showed higher levels of *α*-smooth muscle actin, indicating that anisotropic stress fields significantly increased activation of fibroblasts into myofibroblasts. Error bars represent the standard error with 9 images from isotropic regions (each of them contains 15-51 cells) and 9 images from anisotropic regions (each of them contains 17-47 cells).

To evaluate whether pre-stressed/pre-strained cell models and other thermoelastic-based classical cell models that do not account for tension anisotropy could explain these results, we implemented several other models computationally. All other models predicted the diametric opposite trend (fig. S16) as these models do not account for the effect of tension anisotropy on cell contractility and cell force generation. Although these models capture the observation that ECM stiffness promotes cytoskeletal tension and cell force generation, they fail to capture the effect of tension anisotropy on cell actomyosin contractility and cell force generation.

Thus, although mechanical tensile forces have been shown to promote activation of fibroblasts (*62*), our work reveals that the directionality of stresses is in fact a central determinant that can override stress magnitude. To further establish this, we studied *α*-smooth muscle actin (*α*-SMA) expression which is an indicator of fibroblast activation (Figs. 6C-6F). As predicted by the model, application of stress in a single direction promoted fibroblast activation, while application of the same stress simultaneously in two perpendicular directions inhibited activation.

Together, feedback between cell protrusions and matrix fibers in the early-stages of cell-matrix interaction enables initially round, small, and inactive fibroblasts to spread, polarize, and contract. Through this mechanical crosstalk, protrusions align and stiffen matrix fibers anisotropically, while the anisotropic mechanical microenvironment induced by matrix fiber alignment in turn stabilizes the protrusions in the direction of alignment. Disruption of this mechanical feedback, either by reducing tension anisotropy or inhibiting protrusions at the early-stages of cell-matrix interaction, inhibits activation of fibroblasts. While previous studies have shown cell activation under tensile forces, our results show that these tensile forces should be in an anisotropic form to maximize cell activation. These results shed light on mechanisms of fibrotic disease, and suggest future therapeutic pathways.

## Acknowledgments

D.S. thankfully acknowledges NIH training grant (5-T32-HL07081-38).

## Author contributions

D. Shakiba, F. Alisafaei, E.L. Elson, and G.M. Genin designed the research. D. Shakiba, L.E. Iannucci, M.D. Davidson, K.M. Pryse, P-h G. Chao, J.A. Burdick, S.P. Lake, E.L. Elson, and G.M. Genin performed and supervised the experiments. F. Alisafaei and V. B. Shenoy developed the theoretical models and performed numerical simulations. F. Alisafaei, D. Shakiba, and G.M. Genin wrote the manuscript and all authors discussed the results and commented on the manuscript.

## Data and materials availability

Data, materials, and code for the main text or the supplementary materials are available upon request.

## Supplementary Materials

Materials and Methods

Supplementary Text

Figures S1-S17

Movie S1

References (*63-73*)

