## Supplementary Information for "Tension anisotropy drives phenotypic transitions of cells via two-way cell-ECM feedback"

**This PDF file includes:**

Materials and Methods
Figs. S1 to S17
Captions for Movie S1

**Other Supplementary Materials for this manuscript include the following:**

Movie S1

### Materials and Methods

Cell culture and 3D collagen matrix formation

Human dermal fibroblasts, derived from adult dermis, were purchased from Lonza (Basel, Switzerland, Catalog #: CC-2511). Cells were first cultured at 37°C with 5% $CO_2$ in DMEM containing 10% fetal bovine serum. Next, cells were detached with 0.05% Trypsin/ EDTA for 5 min and separated with Accutase (Sigma Aldrich, St. Louis, MO) then resuspended in fresh DMEM. Finally, the suspended cells in DMEM were mixed with type I collagen in 0.1% acetic acid derived from rat tails and brought to neutral pH by addition of sodium hydroxide to make the final gel concentration 1.0 mg/ml.

For immunostaining of cells, 300 µl of the gel was poured into a 4-well chamber glass-bottom dish (#1.5 glass 0.17-0.19 mm) or a Cellvis 8-well cover glass chamber (#1.5 cover glass, 0.170 mm). Gels were polymerized at 37°C in a tissue culture incubator (5% $CO_2$) for 20-30 minutes. Collagen intensity measurements were performed on gels without the cells to verify that the collagen gels were completely polymerized after 20-30 minutes.

Pre-treatment of resuspended fibroblasts was done with DMEM containing nocodazole (Sigma Aldrich, St Louis) for 30 minutes before making the gel at 37°C with 5% $CO_2$. This ensured that our experiments captured the effect of the inhibitor (nocodazole) at the earliest stage of tissue remodeling. After the first hour of incubation, chambers were filled with DMEM containing nocodazole, and the media was changed every 4 hours to keep the effect of nocodazole on the fibroblast cells.

Quantification of collagen density

We calculated the collagen density as a function of distance from the cell (Figure 1E) from confocal microscopy stacks using a custom routine written in MATLAB (MathWorks, Natick, MA). Image stacks were collected with collagen in the green channel (reflectance) and cells in the red channel (fluorescence). We processed each image plane to define the cell, and calculated the intensity of the collagen signal from the green channel in several concentric, dilated regions surrounding the cell. The laser intensity was set so that the densest regions of collagen would not saturate the reflectance signal, and all other intensities were measured relative to this. The red channels were converted to binary images by thresholding, small spots (less than 20 pixels) were removed by area opening, and internal holes in the cells were filled to produce the final cell image. This image was dilated and the original image subtracted to define a 3D region around the cell. These regions were then applied as masks to the collagen (reflectance) image stack and the intensity in this region was summed and then divided by the region's area to calculate the normalized collagen intensity in the region. This process was repeated several times with increasingly larger dilation values until the final dilated region was close to the image boundaries.

Time-lapse microscopy (live-cell microscopy)

Simultaneous fluorescence confocal imaging of cells labeled with Cell Tracker Orange (Thermo Fisher, Waltham, MA) and confocal reflectance imaging of collagen in 3D polymerized collagen samples were done on a Zeiss LSM 510 microscope using a 40X, 1.2 NA water immersion objective. To ensure that imaged cells experienced 3D microenvironments, cells far from the bottom of the dish were selected and imaged with their surrounding collagen matrix in 3D stacks. The imaging of each cell was repeated every 20 minutes for 2-3 days.

Immunofluorescence microscopy of fibroblasts in 3D collagen gels
To visualize the cytoskeleton organization, 24-hour-old samples were fixed for 20 minutes at room temperature with 4% paraformaldehyde pre-heated to 37°C in DPBS, permeabilized with 0.5% Triton X-100 in DPBS for 10 min, and blocked with 10% normal goat serum (Life Technologies, Carlsbad, CA) for 30 minutes. To stain for tubulin, monoclonal anti-B-tubulin-FITC (1:25 dilution) was added to samples and incubated for 90 minutes at room temperature. To stain for actin, samples were incubated with rhodamine-conjugated phalloidin (1:100 dilution) for 1 hour at room temperature followed by three washes with DPBS/Tween and three washes with DPBS. After washing in DPBS/Tween and DPBS, observations were made using a Zeiss (Oberkochen, Germany) LSM 880 confocal microscope with Airyscan detection, and image stacks were collected.

Strain mapping
We used a strain mapping technique, called Direct Deformation Estimation (DDE) (*53*), to determine the cell-generated strain distribution around the fibroblasts. Briefly, a warping function was estimated that mapped between 3D image stacks taken at the first time point and a 3D image stack taken at a later timepoint. The warping function was designed to provide an estimate of the deformation gradient tensor over defined, overlapping regions within the image volumes. The warping function for each region was optimized using a modification of the Lucas-Kanade algorithm (see reference (*54*)). Using the Zeiss 510 LSM confocal microscope, z-stack images obtained at different time points of development were used as the inputs to the MATLAB code of DDE. To evaluate the strain distribution at each time point, images from two successive time points were utilized during each evaluation. The Green-Lagrange strain tensor was then calculated from the deformation gradient estimated for each region, and by assembling all of the strain regions together, a map of the spatially varying strain distribution around the cell was obtained. The method resolution is limited by the feature size in the 3D texture that is tracked from image to image, in this case, the matrix pore sizes. However, studies on phantom images in 2D and 3D show that accuracy is increased by tracking overlapping regions that each include multiple trackable features (*53*). In this case, regions on the order of 25x25x5 pixels in the 512x512 image stacks were tracked.

Fabrication of aligned collagen gels
2D and 3D collagen matrices with aligned fibers were created using a polydimethylsiloxane (PDMS) (Sylgard 184, Dow Corning) microfluidic device. The details of the microfluidic device are described in (*41*). The liquid level difference between the inlet and the outlet of the device generates a hydrostatic pressure and subsequently a laminar flow which in turn induces collagen alignment due to shear stresses between collagen fibers (Figure S4). To prevent the adhesion of collagen fibers to PDMS, the channel was treated with 3% Pluronic F68 (Sigma) and was sterilized with ethyl alcohol overnight. To enclose the microchannel, silanized glass slides were attached to the channels. Before attachment, the glass slides were first immersed in 5% glutaraldehyde (Electron Microscopy Science) for 30 minutes, rinsed with deionized water, and then immersed in ethyl alcohol for sterilization.

Electrospinning of isotropic and aligned synthetic fibrous hydrogels
Norbornene modified hyaluronic acid (NorHA) was used to make synthetic hydrogel fibers that are cytocompatible and have tunable mechanical and structural properties to mimic natural ECMs (*63, 64*). NorHA was synthesized as previously described (*65*). Briefly, HA was first converted to

the tetrabutylammonium salt form to yield HA that is soluble in dimethyl sulfoxide (DMSO). Sodium HA (66-99 KDa, Lifecore) was dissolved in DI $H_2O$ at 2 wt % with Dowex® resin (50Wx8 ion-change resin, MilliporeSigma) at a ratio of 3:1 (resin: HA by weight) and mixed for 30 minutes at room temperature. The resin was filtered, the filtrate was titrated to pH 7.03 with 1 M tetrabutylammonium hydroxide (TBA-OH), and then frozen and lyophilized. The TBA modification of HA was confirmed with $^1$H NMR. HA-TBA was dissolved in anhydrous DMSO at 2 wt% with 4-(dimethylamino)pyridine (DMAP) (1.5 molar ratio to HA-TBA repeat units) and 5-norbornene-2-carboxylic acid (3:1 molar ratio to HA-TBA repeat units) under a nitrogen atmosphere. Once dissolved, di-tert-butyl dicarbonate ($Boc_2O$, 0.4 M ratio to HA-TBA units) was injected into the vessel and the reaction was carried out for ~20 hrs at 45 °C. The reaction was quenched with 2X cold (4°C) DI $H_2O$ and dialyzed against water with 0.25g NaCl/ L of DI H2O for 3 days. NorHA was then mixed with NaCl (1g NaCl/ 100ml solution) and precipitated in ice-cold acetone (1L acetone/100ml solution). The precipitate was dissolved in DI $H_2O$, dialyzed for 5 days, frozen, and lyophilized. Norbornene functionalization, ~17% modification, was confirmed with $^1$H NMR.

To electrospin NorHA, a 3.5 wt% NorHA, 2.5 wt% PEO (900 kDa, MilliporeSigma), 0.05 (v/v) % Irgacure 2959 (I2959, MilliporeSigma), and 4 mg/ml fluorescent dextran (70 kDa FITC-dextran, MilliporeSigma) were mixed with stoichiometric ratios of dithiothreitol (DTT) to norbornene groups (i.e., 0.2 thiols:norbornenes) in DI $H_2O$, mixed at 250 rpm for 24 hours protected from light, and loaded into syringes for electrospinning. To electrospin NorHA fibers, an 18G blunt tip needle was used with a flow rate of 0.7 ml/hr and a needle to collector distance of 19 cm. A positive voltage (+28-30 kV) was applied to the needle and a negative voltage (-5 kV) was applied to a rotating mandrel, where the speed was adjusted to yield isotropic (slow rotation) or highly aligned (fast rotation) fibrous scaffolds. After electrospinning 150 µl of polymer solution, the fibrous scaffold was removed from the mandrel, purged with nitrogen, and photocrosslinked with UV light (10 mW/cm$^2$, 320-395 nm, Omnicure S2000) for 1 hour on each side of the scaffold.

For cell culture studies, 2 cm x 2 cm sections of scaffolds were sterilized in a cell culture hood with germicidal UV for 1 hour, hydrated in DPBS, and then conjugated with cell adhesive peptides containing the RGD sequence (sequence: GCGYGRGDSPG, GenScript) as previously described (*64*). Briefly, a solution containing 1 mM RGD peptide and 0.05 (v/v) % I2959 diluted in DPBS was applied to scaffolds, and thiols on cystines within the peptide were conjugated to norbornene groups on the fibers through a thiol-ene photoinitiated reaction (5 mW/ cm$^2$ UV, 5 minutes). Scaffolds were washed 3 times with DPBS, and then cells were seeded on scaffolds at a concentration of $1.5 \times 10^5$ cells/ml. Cells were cultured on isotropic and aligned scaffolds for various times, fixed with formalin, and stained with Alexa Fluor 647 Phalloidin conjugate (1:400, ThermoFisher Scientific).

Cruciform tissue analog fabrication

Fibroblast-seeded collagen hydrogel tissue analogs (TAs) were fabricated using previously established methods (*66*). Collagen was acid-extracted from Long Evans rat tail tendons and stored in 0.1% acetic acid at a stock concentration of 2.25 mg/mL at 4°C (*67*). Neonatal human dermal fibroblasts (nHDFs; ATCC PCS-201-010) were expanded out to passage 5 in complete culture medium (DMEM, 10% FBS, 1% Pen/Strep). Stock collagen solution was neutralized to a final pH of 7.0 and 300 mOsm through addition of 1 N NaOH, 10x PBS, and 1x PBS and mixed with isolated nHDFs for a final collagen concentration of 1.5 mg/mL and cell concentration of 5 x 105 cells/mL. Cell-collagen mixture was then injected into cruciform-shaped Teflon mold. The molds contained a textured borosilicate rod at the end of each to encourage directed fiber formation. After injection-molding, the TAs were placed in an incubator at 37°C for 30 minutes before being covered with 5 mL complete culture media and subsequently static cultured for 5 days. Media was changed every 2-3 days.

Three-dimensional cytoskeletal model

The cell model is composed of the cytoskeleton, the nucleus, and the focal adhesion (*55*). In this section, we describe the cytoskeleton which includes (i) the myosin molecular motors, (ii) the actin filaments, and (iii) the microtubules.

**1. Myosin motors, actin filaments, and microtubules**

The first component of the cytoskeletal model is myosin which generates cell internal contractility and is denoted by a contractility tensor, $\rho_{ij}$, whose components represent cell internally-generated stresses in different directions (*56*). In what follows, we first define the contractility tensor $\rho_{ij}$ and we then discuss how the cell model responds to mechanical signals from their extracellular environment by adjusting $\rho_{ij}$.

The force generated by a myosin motor at submicron levels can be treated as a force dipole (Figure S17). Force dipoles are pairs of equal but oppositely directed forces $F_i(x_j)$ and $-F_i(x_j+\Delta x_j)$ where $|\Delta x_j|$ is the length of myosin II filaments (approximately 200 nm) (*68*), and $x_j$ and $x_j+\Delta x_j$ are coordinates of myosin head domains. We then can define the work done by a force dipole $W_{\text{dipole}}$ and the total work generated by all myosin motors per volume $W$ as follows, respectively,

$$W_{\text{dipole}} = F_i u_i(x_j+\Delta x_j) - F_i u_i(x_j) \tag{S1.1}$$

$$W = \left(\frac{1}{V}\right)\sum_{k=1}^{N}\left[F_i^{(k)} u_i\left(x_j^{(k)}+\Delta x_j^{(k)}\right) - F_i^{(k)} u_i\left(x_j^{(k)}\right)\right] \tag{S1.2}$$

where $u_i(x_j)$ and $u_i(x_j+\Delta x_j)$ are the displacements of the cytoskeleton at $x_j$ and $x_j+\Delta x_j$, respectively, and $V$ is the volume of the cytoplasm. Assuming that $N$ is the total number of bound (phosphorylated) myosin motors and using equation (S1.4), we can rewrite equation (S1.2) in this form

$$W = \left(\frac{1}{V}\right)\sum_{k=1}^{N} F_i^{(k)}\Delta x_j^{(k)}\partial_j u_i \tag{S1.3}$$

$$\partial_j u_i = \frac{u_i\left(x_j^{(k)}+\Delta x_j^{(k)}\right) - u_i\left(x_j^{(k)}\right)}{\Delta x_j^{(k)}} \tag{S1.4}$$

Therefore, the total work can be presented as a function of the contractility tensor $\rho_{ij}$

$$W = \rho_{ij}\varepsilon_{ij} \tag{S1.5}$$

where

$$\varepsilon_{ij} = \frac{1}{2}\left(\partial_j u_i + \partial_i u_j\right) \tag{S1.6}$$

is the linearized strain, and

$$\rho_{ij} = \left(\frac{1}{V}\right)\sum_{k=1}^{N} F_i^{(k)} \Delta x_j^{(k)} \tag{S1.7}$$

Note that the contractility tensor $\rho_{ij}$ is a symmetric tensor and holds the following condition

$$F_i^{(k)} \Delta x_j^{(k)} = F_j^{(k)} \Delta x_i^{(k)} \tag{S1.8}$$

Equation (S1.7) demonstrates that the cell contractility tensor $\rho_{ij}$ represents the density of phosphorylated myosin molecular motors. Experimental observations, however, have shown that cells adjust the density of phosphorylated myosin in response to mechanical signals from the environment. For example, cells on/within stiff matrices have higher densities of phosphorylated myosin and are more contractile than those on/within soft matrices (*69*, *70*). Therefore, to have a contractile model that actively responds to extracellular mechanical signals, we hypothesize that the average of contractility in all three directions, $\frac{1}{3}\rho_{kk} = (\rho_{11} + \rho_{22} + \rho_{33})/3$, in our coarse-grain model increases with the average of tension in the actin filament network, $\frac{1}{3}\sigma_{kk} = (\sigma_{11} + \sigma_{22} + \sigma_{33})/3$

$$\frac{\rho_{kk}}{3} = f_{\mathrm{m}}\frac{\sigma_{kk}}{3} + f_0\,\rho_0 \tag{S1.9}$$

where this tension-dependent feedback mechanism is regulated by the feedback constant parameter $f_{\mathrm{m}}$. In equation (S1.9), the constant parameter $\rho_0$ is the initial contractility (basal cell contractility), and the constant parameter $f_0$ regulates the mean contractility $\frac{1}{3}\rho_{kk}$ in the absence of tension ($\sigma_{kk} = 0$). We have previously shown that as a result of the feedback mechanism in (S1.9), in addition to the cell contractility $\rho_{ij}$, the stiffness of the actin network $C_{ijkl}^{(\mathrm{A})}$ and the tension it carries, $\sigma_{ij}$, also increase with matrix stiffness in an orientation-dependent manner.

To be able to implement the model in (S1.9) into a three-dimensional finite element model, we need to derive the total stress $\sigma_{ij}$ and total stiffness $C_{ij}$ tensors. To this end, we start with the following definition for $\rho_{ij}$ which relates it to the strain tensor $\boldsymbol{\varepsilon}^{(\mathrm{X})}$ (the three-dimensional representation of $\varepsilon^{(\mathrm{X})}$ shown in Figure S17)

$$\rho_{ij} = K^{(\rho)}\varepsilon_{kk}^{(\mathrm{X})}\delta_{ij} + 2\mu^{(\rho)}\left(\varepsilon_{ij}^{(\mathrm{X})} - \frac{1}{3}\varepsilon_{kk}^{(\mathrm{X})}\delta_{ij}\right) + \bar{\rho}_0\delta_{ij} \tag{S1.10}$$

where

$$K^{(\rho)} = \frac{3K^{(\mathrm{MT})}\alpha_{\mathrm{v}} - 1}{3(\beta_{\mathrm{v}} - \alpha_{\mathrm{v}})} \tag{S1.11}$$

is effective modulus for the motor density,

$$\mu^{(\rho)} = \frac{2\mu^{(\mathrm{MT})}\alpha_{\mathrm{d}} - 1}{2(\beta_{\mathrm{d}} - \alpha_{\mathrm{d}})} \tag{S1.12}$$

is the effective modulus of the polarization,

$$\bar{\rho}_0 = \frac{\beta_{\mathrm{v}}\,\rho_0}{\beta_{\mathrm{v}} - \alpha_{\mathrm{v}}} \tag{S1.13}$$

is the effective contractility,

$$K^{(\mathrm{MT})} = \frac{E^{(\mathrm{MT})}}{3(1 - 2v^{(\mathrm{MT})})} \tag{S1.14}$$

is the bulk modulus of the cytoskeletal components that are in compression (*e.g.*, the microtubule network), and

$$\mu^{(\mathrm{MT})} = \frac{E^{(\mathrm{MT})}}{2(1 + v^{(\mathrm{MT})})} \tag{S1.15}$$

is the shear modulus of the cytoskeletal components that are in compression (*e.g.*, the microtubule network). $E^{(\mathrm{MT})}$ in these equations is the elastic modulus of the cytoskeletal components that are in compression (*e.g.*, the microtubule network), $v^{(\mathrm{MT})}$ is the Poisson's ratio of the cytoskeletal components that are in compression (*e.g.*, the microtubule network), $\rho_0$ is the initial cell contractility, $\alpha_{\mathrm{v}}$ is the volumetric chemo-mechanical feedback parameter (large values of $\alpha_{\mathrm{v}}$ lead to higher phosphorylation of myosin), $\alpha_{\mathrm{d}}$ is the deviatoric chemo-mechanical feedback parameter which represents the cell tendency to polarized contraction (small values of $\alpha_{\mathrm{d}}$ results in non-polarized contractility), $\beta_{\mathrm{v}}$ is the volumetric chemical stiffness parameter which regulates the mean contractility $\frac{1}{3}\rho_{kk}$ in the absence of tension ($\sigma_{kk} = 0$) by maintaining the contractility at a basal level (large values of $\beta_{\mathrm{v}}$ keeps the level of myosin phosphorylation low), and $\beta_{\mathrm{d}}$ is the deviatoric chemical stiffness parameter which represents the disinclination of myosin to orient along the cell's polarization direction (large values of $\beta_{\mathrm{d}}$ leads to a random orientation of myosin motors).
A large portion of microtubules experiences compression due to the cell's internal contractile forces (*71, 72*). Consistently, the cell contractility $\rho_{ij}$ in our model generates the compressive stress $C^{(\mathrm{MT})}_{ijkl}\varepsilon^{(\mathrm{X})}_{kl}$ on microtubules where

$$C^{(\mathrm{MT})}_{ijkl} = K^{(\mathrm{MT})}\delta_{ij}\delta_{kl} + \mu^{(\mathrm{MT})}\left(\delta_{ik}\delta_{jk} + \delta_{il}\delta_{jk} - \frac{2}{3}\delta_{ij}\delta_{kl}\right) \tag{S1.16}$$

is the microtubule network stiffness tensor. Unlike microtubules, actin filaments experience tension. Similarly, the cell contractility $\rho_{ij}$ in our model generates tension in the actin network, $\sigma_{ij}$, in addition to compressing microtubules,

$$\rho_{ij} = -C^{(\mathrm{MT})}_{ijkl}\varepsilon^{(\mathrm{X})}_{kl} + \sigma_{ij} \tag{S1.17}$$

In our model, Equation (S1.10) indicates that the contractility tensor $\rho_{ij}$ is initially isotropic and, as a result, the cell exhibits the same contractility in all directions in the initial configuration. This can be mathematically shown by rewriting equation (S1.10) in the following form

$$\rho_{ij} = C^{(\rho)}_{ijkl}\varepsilon^{(X)}_{kl} + \bar{\rho}_0\delta_{ij} \tag{S1.18}$$

where

$$C^{(\rho)}_{ijkl} = K^{(\rho)}\delta_{ij}\delta_{kl} + \mu^{(\rho)}\left(\delta_{ik}\delta_{jk} + \delta_{il}\delta_{jk} - \frac{2}{3}\delta_{ij}\delta_{kl}\right) \tag{S1.19}$$

As shown in Figure S13, the model predicts that the contractility can become nonuniform and anisotropic, dependent on extracellular physical constraints. However, for all simulations, we start with a uniform (independent of spatial location) and isotropic (independent of direction) contractility. Equations (S1.17) and (S1.18) demonstrate that, in the stress-free condition $\sigma_{ij} = 0$ (initially there is no tension), the diagonal components of $\rho_{ij}$ are all equal and non-zero $\rho_{11} = \rho_{22} = \rho_{33} \neq 0$, while the off-diagonal components are all zero $\rho_{12} = \rho_{21} = \rho_{13} = \rho_{31} = \rho_{23} = \rho_{32} = 0$, indicating that the contractility tensor $\rho_{ij}$ is initially isotropic. This is better shown by writing equation (S1.18) in the following format using Voigt notation

$$\begin{Bmatrix} \rho_{11} \\ \rho_{22} \\ \rho_{33} \\ \rho_{12} \\ \rho_{13} \\ \rho_{23} \end{Bmatrix} = \begin{bmatrix} C^{(\rho)}_{1111} & C^{(\rho)}_{1122} & C^{(\rho)}_{1133} & C^{(\rho)}_{1112} & C^{(\rho)}_{1113} & C^{(\rho)}_{1123} \\ C^{(\rho)}_{2211} & C^{(\rho)}_{2222} & C^{(\rho)}_{2233} & C^{(\rho)}_{2212} & C^{(\rho)}_{2213} & C^{(\rho)}_{2223} \\ C^{(\rho)}_{3311} & C^{(\rho)}_{3322} & C^{(\rho)}_{3333} & C^{(\rho)}_{3312} & C^{(\rho)}_{3313} & C^{(\rho)}_{3323} \\ C^{(\rho)}_{1211} & C^{(\rho)}_{1222} & C^{(\rho)}_{1233} & C^{(\rho)}_{1212} & C^{(\rho)}_{1213} & C^{(\rho)}_{1223} \\ C^{(\rho)}_{1311} & C^{(\rho)}_{1322} & C^{(\rho)}_{1333} & C^{(\rho)}_{1312} & C^{(\rho)}_{1313} & C^{(\rho)}_{1323} \\ C^{(\rho)}_{2311} & C^{(\rho)}_{2322} & C^{(\rho)}_{2333} & C^{(\rho)}_{2312} & C^{(\rho)}_{2313} & C^{(\rho)}_{2323} \end{bmatrix} \begin{Bmatrix} \varepsilon_{11} \\ \varepsilon_{22} \\ \varepsilon_{33} \\ \varepsilon_{12} \\ \varepsilon_{13} \\ \varepsilon_{23} \end{Bmatrix} + \begin{Bmatrix} \bar{\rho}_0 \\ \bar{\rho}_0 \\ \bar{\rho}_0 \\ 0 \\ 0 \\ 0 \end{Bmatrix} \tag{S1.20}$$

where the first, second, and third components of the last term in equation (S1.20) have the same value $\bar{\rho}_0$.

Concomitant with phosphorylation of more myosin, cells also respond to tension by polymerizing actin filaments and bundling and aligning them in the direction of tension (*32*, *73*). Thus, in addition to the feedback mechanism between the contractility $\rho_{ij}$ and the stress $\sigma_{ij}$ in (S1.9), we also hypothesize that the stiffness of the actin network $C^{(A)}_{ijkl}$ increases in proportion to, and in the directions of, the tensile principal components of the stress tensor $\sigma_{ij}$, (*55*)

$$C^{(A)}_{ijkl} = C^{(I)}_{ijkl} + C^{(F)}_{ijkl} \tag{S1.21}$$

$\mathbf{C}^{(F)}$ denotes the stiffening of the actin network with tension (but not in compression) and $\mathbf{C}^{(I)}$ is the initial stiffness of the actin filaments network

$$C^{(I)}_{ijkl} = K^{(I)}\delta_{ij}\delta_{kl} + \mu^{(I)}\left(\delta_{ik}\delta_{jk} + \delta_{il}\delta_{jk} - \frac{2}{3}\delta_{ij}\delta_{kl}\right) \tag{S1.22}$$

where

$$K^{(\mathrm{I})} = \frac{E^{(\mathrm{I})}}{3(1 - 2v^{(\mathrm{I})})} \tag{S1.23}$$

is the initial bulk modulus of the actin network, and

$$\mu^{(\mathrm{I})} = \frac{E^{(\mathrm{I})}}{2(1 + v^{(\mathrm{I})})} \tag{S1.24}$$

is the initial shear modulus of the actin network, $E^{(\mathrm{I})}$ is the initial elastic modulus of the actin network, and $v^{(\mathrm{I})}$ is the initial Poisson's ratio of the actin network.

To define the stiffening part of the stiffness tensor, $\mathbf{C}^{(\mathrm{F})}$, we first decompose $\boldsymbol{\sigma}$

$$\sigma_{ij} = \sigma_{ij}^{(\mathrm{A})} = \sigma_{ij}^{(\mathrm{I})} + \sigma_{ij}^{(\mathrm{F})} \tag{S1.25}$$

where $\boldsymbol{\sigma}^{(\mathrm{I})}$

$$\sigma_{ij}^{(\mathrm{I})} = \mathrm{C}_{ijkl}^{(\mathrm{I})}\, \varepsilon_{kl}^{(\mathrm{Y})} \tag{S1.26}$$

is linearly related to the strain tensor $\boldsymbol{\varepsilon}^{(\mathrm{Y})}$ (the three-dimensional representation of $\varepsilon^{(\mathrm{Y})}$ shown in Figure S17) which can be written as a function of its eigenvalues (principal strains) $\varepsilon_1^{(\mathrm{Y})}, \varepsilon_2^{(\mathrm{Y})}, \varepsilon_3^{(\mathrm{Y})}$ and eigenvectors $\boldsymbol{n}_1, \boldsymbol{n}_2, \boldsymbol{n}_3$

$$\boldsymbol{\varepsilon}^{(\mathrm{Y})} = \sum_{i=1}^{3} \varepsilon_i^{(\mathrm{Y})} \boldsymbol{n}_i \otimes \boldsymbol{n}_i = \sum_{i=1}^{3} \varepsilon_i^{(\mathrm{Y})} \mathbf{E}_i \tag{S1.27}$$

where the symmetric tensors $\mathbf{E}_1 = \boldsymbol{n}_1 \otimes \boldsymbol{n}_1$, $\mathbf{E}_2 = \boldsymbol{n}_2 \otimes \boldsymbol{n}_2$, and $\mathbf{E}_3 = \boldsymbol{n}_3 \otimes \boldsymbol{n}_3$ are the eigenprojections of $\boldsymbol{\varepsilon}^{(\mathrm{Y})}$ and $\otimes$ denotes the dyadic product of two arbitrary vectors $\boldsymbol{u}$ and $\boldsymbol{v}$ as $(\boldsymbol{u} \otimes \boldsymbol{v})_{ij} = u_i v_j$. With the eigenvalues ($\varepsilon_1^{(\mathrm{Y})}$, $\varepsilon_2^{(\mathrm{Y})}$, $\varepsilon_3^{(\mathrm{Y})}$) and eigenvectors ($\boldsymbol{n}_1$, $\boldsymbol{n}_2$, $\boldsymbol{n}_3$) at hand, we next define $\sigma_{ij}^{(\mathrm{F})}$ in (S1.25)

$$\boldsymbol{\sigma}^{(\mathrm{F})} = \sum_{i=1}^{3} \frac{\partial f(\varepsilon_i^{(\mathrm{Y})})}{\partial \varepsilon_i^{(\mathrm{Y})}} \boldsymbol{n}_i \otimes \boldsymbol{n}_i = \sum_{i=1}^{3} \sigma^{(\mathrm{F})}\left(\varepsilon_i^{(\mathrm{Y})}\right) \mathbf{E}_i = \sum_{i=1}^{3} \sigma_i^{(\mathrm{F})} \mathbf{E}_i \tag{S1.28}$$

where $\sigma_i^{(\mathrm{F})}$ are the eigenvalues (principal stresses) of the stress tensor $\sigma_{ij}^{(\mathrm{F})}$ and are defined as follows

$$\sigma_i^{(\mathrm{F})} = \frac{\partial f(\varepsilon_i^{(\mathrm{Y})})}{\partial \varepsilon_i^{(\mathrm{Y})}} = \tag{S1.29}$$

$$\begin{cases} 0 & \varepsilon_i^{(\mathrm{Y})} < \epsilon_1 \\ \ell \dfrac{\left(\dfrac{\varepsilon_i^{(\mathrm{Y})} - \epsilon_1}{\epsilon_2 - \epsilon_1}\right)^t \left(\varepsilon_i^{(\mathrm{Y})} - \epsilon_1\right)^2}{(t+1)(t+2)} & \epsilon_1 \le \varepsilon_i^{(\mathrm{Y})} < \epsilon_2 \\ \ell \left[\dfrac{\left(1 + \varepsilon_i^{(\mathrm{Y})} - \epsilon_2\right)^{s+2} - 1}{(s+1)(s+2)} + \dfrac{\epsilon_2 - \varepsilon_i^{(\mathrm{Y})}}{s+1} + \dfrac{(\varepsilon_i^{(\mathrm{Y})} - \epsilon_2)(\epsilon_2 - \epsilon_1)}{t+1} + \dfrac{(\epsilon_2 - \epsilon_1)^2}{(t+1)(t+2)}\right] & \varepsilon_i^{(\mathrm{Y})} \ge \epsilon_2 \end{cases}$$

to ensure the continuity and smoothness of the first and second derivatives of $\sigma_i^{(\mathrm{F})}$ with respect to $\varepsilon_i^{(\mathrm{Y})}$ at the transition points $\epsilon_1 = \epsilon_\mathrm{c} - 0.5\epsilon_\mathrm{t}$ and $\epsilon_2 = \epsilon_\mathrm{c} + 0.5\epsilon_\mathrm{t}$ where $\epsilon_\mathrm{t} = 0.25\epsilon_\mathrm{c}$ is the transition width, $\epsilon_\mathrm{c}$ is the critical (tensile) principal strain, and $t$ is the transition constant. Equation (S1.29) shows that for large tensile strains $\varepsilon_i^{(\mathrm{Y})} \ge \epsilon_2$, the principal stress $\sigma_i^{(\mathrm{F})}$ nonlinearly increases with the principal strain $\varepsilon_i^{(\mathrm{Y})}$ where this increase is regulated by the stiffening parameters $\ell$ and $s$. With $\boldsymbol{\sigma}^{(\mathrm{F})}$ at hand from equation (S1.28), we can now determine $\mathbf{C}^{(\mathrm{F})}$ as follows

$$\mathrm{C}_{ijkl}^{(\mathrm{F})} = \frac{\mathrm{d}\sigma_{ij}^{(\mathrm{F})}}{\mathrm{d}\varepsilon_{kl}^{(\mathrm{Y})}} \qquad \text{or} \qquad \mathbf{C}^{(\mathrm{F})} = \frac{\mathrm{d}\boldsymbol{\sigma}^{(\mathrm{F})}}{\mathrm{d}\boldsymbol{\varepsilon}^{(\mathrm{Y})}} \tag{S1.30}$$

The piecewise linear approximation in equation (S1.30) requires $\sigma_{ij}^{(\mathrm{F})}$ as a function of $\varepsilon_{ij}^{(\mathrm{Y})}$ while equation (S1.29) gives $\sigma_i^{(\mathrm{F})}$ as a function of $\varepsilon_i^{(\mathrm{Y})}$. Thus, we first write $\mathrm{C}_{ijkl}^{(\mathrm{F})}$ in the following form using the definition of $\sigma_{ij}^{(\mathrm{F})}$ in equation (S1.28)

$$\mathbf{C}^{(\mathrm{F})} = \sum_{i=1}^{3} \left\{ \mathbf{E}_i \otimes \frac{\mathrm{d}\sigma_i^{(\mathrm{F})}}{\mathrm{d}\boldsymbol{\varepsilon}^{(\mathrm{Y})}} + \sigma_i^{(\mathrm{F})} \frac{\mathrm{d}\mathbf{E}_i}{\mathrm{d}\boldsymbol{\varepsilon}^{(\mathrm{Y})}} \right\} \tag{S1.31}$$

and we then expand the first term in the right-hand side of equation (S1.31) by applying the chain rule

$$\mathbf{C}^{(\mathrm{F})} = \sum_{i=1}^{3} \left\{ \sum_{j=1}^{3} \frac{\partial \sigma_i^{(\mathrm{F})}}{\partial \varepsilon_j^{(\mathrm{Y})}} \mathbf{E}_i \otimes \frac{\mathrm{d}\varepsilon_j^{(\mathrm{Y})}}{\mathrm{d}\boldsymbol{\varepsilon}^{(\mathrm{Y})}} + \sigma_i^{(\mathrm{F})} \frac{\mathrm{d}\mathbf{E}_i}{\mathrm{d}\boldsymbol{\varepsilon}^{(\mathrm{Y})}} \right\} \tag{S1.32}$$

To further expand equation (1.32) and derive an exact form for $\mathbf{C}^{(\mathrm{F})}$, we consider the three following cases.

In the first case, the three eigenvalues of the strain tensor $\varepsilon_{ij}^{(\mathrm{Y})}$ are all nonidentical ($\varepsilon_1^{(\mathrm{Y})} \neq \varepsilon_2^{(\mathrm{Y})} \neq \varepsilon_3^{(\mathrm{Y})}$). In this case, we can derive the following expression for $\mathbf{C}^{(\mathrm{F})}$ from equation (S1.32) by taking the derivatives of $\varepsilon_j^{(\mathrm{Y})}$ and $\mathbf{E}_i$ with respect to $\boldsymbol{\varepsilon}^{(\mathrm{Y})}$

$$\mathbf{C}^{(\mathrm{F})} = \sum_{a=1}^{3} \frac{\sigma_a^{(\mathrm{F})}}{(\varepsilon_a^{(\mathrm{Y})} - \varepsilon_b^{(\mathrm{Y})})(\varepsilon_a^{(\mathrm{Y})} - \varepsilon_c^{(\mathrm{Y})})} \left\{ \frac{\mathrm{d}\left(\boldsymbol{\varepsilon}^{(\mathrm{Y})}\right)^2}{\mathrm{d}\boldsymbol{\varepsilon}^{(\mathrm{Y})}} - \left(\varepsilon_b^{(\mathrm{Y})} + \varepsilon_c^{(\mathrm{Y})}\right)\mathbf{I}_\mathrm{S} - \left[\left(\varepsilon_a^{(\mathrm{Y})} - \varepsilon_b^{(\mathrm{Y})}\right) + \left(\varepsilon_a^{(\mathrm{Y})} - \varepsilon_c^{(\mathrm{Y})}\right)\right] \mathbf{E}_a \otimes \mathbf{E}_a - \left(\varepsilon_b^{(\mathrm{Y})} - \varepsilon_c^{(\mathrm{Y})}\right)(\mathbf{E}_b \otimes \mathbf{E}_b - \mathbf{E}_c \otimes \mathbf{E}_c) \right\} + \sum_{i=1}^{3}\sum_{j=1}^{3} \frac{\partial \sigma_i^{(\mathrm{F})}}{\partial \varepsilon_j^{(\mathrm{Y})}} \mathbf{E}_i \otimes \mathbf{E}_j \qquad \text{(S1.33)}$$

where

$$\left(\frac{\mathrm{d}\left(\boldsymbol{\varepsilon}^{(\mathrm{Y})}\right)^2}{\mathrm{d}\boldsymbol{\varepsilon}^{(\mathrm{Y})}}\right)_{ijkl} = \frac{1}{2}\left(\delta_{ik}\varepsilon_{lj}^{(\mathrm{Y})} + \delta_{il}\varepsilon_{kj}^{(\mathrm{Y})} + \delta_{jl}\varepsilon_{ik}^{(\mathrm{Y})} + \delta_{kj}\varepsilon_{il}^{(\mathrm{Y})}\right) \qquad \text{(S1.34)}$$

is the derivative of the square of $\boldsymbol{\varepsilon}^{(\mathrm{Y})}$, and

$$(\mathbf{I}_\mathrm{S})_{ijkl} = \frac{1}{2}\left(\delta_{ik}\delta_{jl} + \delta_{il}\delta_{jk}\right) \qquad \text{(S1.35)}$$

is the symmetric identity tensor.

In the second case, $\varepsilon_{ij}^{(\mathrm{Y})}$ has two identical eigenvalues ($\varepsilon_1^{(\mathrm{Y})} \neq \varepsilon_2^{(\mathrm{Y})} = \varepsilon_3^{(\mathrm{Y})}$) which gives the following analytical expression for $\mathbf{C}^{(\mathrm{F})}$

$$\mathbf{C}^{(\mathrm{F})} = s_1 \frac{\mathrm{d}\left(\boldsymbol{\varepsilon}^{(\mathrm{Y})}\right)^2}{\mathrm{d}\boldsymbol{\varepsilon}^{(\mathrm{Y})}} - s_2\mathbf{I}_\mathrm{S} - s_3\boldsymbol{\varepsilon}^{(\mathrm{Y})} \otimes \boldsymbol{\varepsilon}^{(\mathrm{Y})} + s_4\boldsymbol{\varepsilon}^{(\mathrm{Y})} \otimes \mathbf{I} + s_5\mathbf{I} \otimes \boldsymbol{\varepsilon}^{(\mathrm{Y})} - s_6\mathbf{I} \otimes \mathbf{I} \qquad \text{(S1.36)}$$

where

$$\mathrm{I}_{ij} = \delta_{ij} \qquad \text{(S1.37)}$$

is the second-order identity tensor, and

$$s_1 = \frac{\sigma_a^{(\mathrm{F})} - \sigma_c^{(\mathrm{F})}}{\left(\varepsilon_a^{(\mathrm{Y})} - \varepsilon_c^{(\mathrm{Y})}\right)^2} + \frac{1}{\varepsilon_a^{(\mathrm{Y})} - \varepsilon_c^{(\mathrm{Y})}}\left(\frac{\partial \sigma_c^{(\mathrm{F})}}{\partial \varepsilon_b^{(\mathrm{Y})}} - \frac{\partial \sigma_c^{(\mathrm{F})}}{\partial \varepsilon_c^{(\mathrm{Y})}}\right) \qquad \text{(S1.38a)}$$

$$s_2 = 2\varepsilon_c^{(\mathrm{Y})} \frac{\sigma_a^{(\mathrm{F})} - \sigma_c^{(\mathrm{F})}}{\left(\varepsilon_a^{(\mathrm{Y})} - \varepsilon_c^{(\mathrm{Y})}\right)^2} + \frac{\varepsilon_a^{(\mathrm{Y})} + \varepsilon_c^{(\mathrm{Y})}}{\varepsilon_a^{(\mathrm{Y})} - \varepsilon_c^{(\mathrm{Y})}}\left(\frac{\partial \sigma_c^{(\mathrm{F})}}{\partial \varepsilon_b^{(\mathrm{Y})}} - \frac{\partial \sigma_c^{(\mathrm{F})}}{\partial \varepsilon_c^{(\mathrm{Y})}}\right) \qquad \text{(S1.38b)}$$

$$s_3 = 2\frac{\sigma_a^{(\mathrm{F})} - \sigma_c^{(\mathrm{F})}}{\left(\varepsilon_a^{(\mathrm{Y})} - \varepsilon_c^{(\mathrm{Y})}\right)^3} + \frac{1}{\left(\varepsilon_a^{(\mathrm{Y})} - \varepsilon_c^{(\mathrm{Y})}\right)^2}\left(\frac{\partial \sigma_a^{(\mathrm{F})}}{\partial \varepsilon_c^{(\mathrm{Y})}} + \frac{\partial \sigma_c^{(\mathrm{F})}}{\partial \varepsilon_a^{(\mathrm{Y})}} - \frac{\partial \sigma_a^{(\mathrm{F})}}{\partial \varepsilon_a^{(\mathrm{Y})}} - \frac{\partial \sigma_c^{(\mathrm{F})}}{\partial \varepsilon_c^{(\mathrm{Y})}}\right) \qquad \text{(S1.38c)}$$

$$s_4 = 2\varepsilon_c^{(\mathrm{Y})}\frac{\sigma_a^{(\mathrm{F})} - \sigma_c^{(\mathrm{F})}}{\left(\varepsilon_a^{(\mathrm{Y})} - \varepsilon_c^{(\mathrm{Y})}\right)^3} + \frac{1}{\varepsilon_a^{(\mathrm{Y})} - \varepsilon_c^{(\mathrm{Y})}}\left(\frac{\partial\sigma_a^{(\mathrm{F})}}{\partial\varepsilon_c^{(\mathrm{Y})}} - \frac{\partial\sigma_c^{(\mathrm{F})}}{\partial\varepsilon_b^{(\mathrm{Y})}}\right)$$
$$+ \frac{\varepsilon_c^{(\mathrm{Y})}}{\left(\varepsilon_a^{(\mathrm{Y})} - \varepsilon_c^{(\mathrm{Y})}\right)^2}\left(\frac{\partial\sigma_a^{(\mathrm{F})}}{\partial\varepsilon_c^{(\mathrm{Y})}} + \frac{\partial\sigma_c^{(\mathrm{F})}}{\partial\varepsilon_a^{(\mathrm{Y})}} - \frac{\partial\sigma_a^{(\mathrm{F})}}{\partial\varepsilon_a^{(\mathrm{Y})}} - \frac{\partial\sigma_c^{(\mathrm{F})}}{\partial\varepsilon_c^{(\mathrm{Y})}}\right) \quad \text{(S1.38d)}$$

$$s_5 = 2\varepsilon_c^{(\mathrm{Y})}\frac{\sigma_a^{(\mathrm{F})} - \sigma_c^{(\mathrm{F})}}{\left(\varepsilon_a^{(\mathrm{Y})} - \varepsilon_c^{(\mathrm{Y})}\right)^3} + \frac{1}{\varepsilon_a^{(\mathrm{Y})} - \varepsilon_c^{(\mathrm{Y})}}\left(\frac{\partial\sigma_c^{(\mathrm{F})}}{\partial\varepsilon_a^{(\mathrm{Y})}} - \frac{\partial\sigma_c^{(\mathrm{F})}}{\partial\varepsilon_b^{(\mathrm{Y})}}\right)$$
$$+ \frac{\varepsilon_c^{(\mathrm{Y})}}{\left(\varepsilon_a^{(\mathrm{Y})} - \varepsilon_c^{(\mathrm{Y})}\right)^2}\left(\frac{\partial\sigma_a^{(\mathrm{F})}}{\partial\varepsilon_c^{(\mathrm{Y})}} + \frac{\partial\sigma_c^{(\mathrm{F})}}{\partial\varepsilon_a^{(\mathrm{Y})}} - \frac{\partial\sigma_a^{(\mathrm{F})}}{\partial\varepsilon_a^{(\mathrm{Y})}} - \frac{\partial\sigma_c^{(\mathrm{F})}}{\partial\varepsilon_c^{(\mathrm{Y})}}\right) \quad \text{(S1.38e)}$$

$$s_6 = 2\varepsilon_c^{(\mathrm{Y})}\frac{\sigma_a^{(\mathrm{F})} - \sigma_c^{(\mathrm{F})}}{\left(\varepsilon_a^{(\mathrm{Y})} - \varepsilon_c^{(\mathrm{Y})}\right)^3} + \frac{\varepsilon_a^{(\mathrm{Y})}\varepsilon_c^{(\mathrm{Y})}}{\left(\varepsilon_a^{(\mathrm{Y})} - \varepsilon_c^{(\mathrm{Y})}\right)^2}\left(\frac{\partial\sigma_a^{(\mathrm{F})}}{\partial\varepsilon_c^{(\mathrm{Y})}} + \frac{\partial\sigma_c^{(\mathrm{F})}}{\partial\varepsilon_a^{(\mathrm{Y})}}\right) - \frac{\left(\varepsilon_c^{(\mathrm{Y})}\right)^2}{\left(\varepsilon_a^{(\mathrm{Y})} - \varepsilon_c^{(\mathrm{Y})}\right)^2}\left(\frac{\partial\sigma_a^{(\mathrm{F})}}{\partial\varepsilon_a^{(\mathrm{Y})}} + \frac{\partial\sigma_c^{(\mathrm{F})}}{\partial\varepsilon_c^{(\mathrm{Y})}}\right)$$
$$- \frac{\varepsilon_a^{(\mathrm{Y})} + \varepsilon_c^{(\mathrm{Y})}}{\varepsilon_a^{(\mathrm{Y})} - \varepsilon_c^{(\mathrm{Y})}}\frac{\partial\sigma_c^{(\mathrm{F})}}{\partial\varepsilon_b^{(\mathrm{Y})}} \quad \text{(S1.38f)}$$

are constants with $(a, b, c)$ being cyclic permutations of (1,2,3).

In the third case, the three eigenvalues of $\varepsilon_{ij}^{(\mathrm{Y})}$ are all identical ($\varepsilon_1^{(\mathrm{Y})} = \varepsilon_2^{(\mathrm{Y})} = \varepsilon_3^{(\mathrm{Y})}$) which gives the following expression for $\mathbf{C}^{(\mathrm{F})}$

$$\mathbf{C}^{(\mathrm{F})} = \left(\frac{\partial\sigma_1^{(\mathrm{F})}}{\partial\varepsilon_1^{(\mathrm{Y})}} - \frac{\partial\sigma_1^{(\mathrm{F})}}{\partial\varepsilon_2^{(\mathrm{Y})}}\right)\mathbf{I}_\mathrm{S} + \frac{\partial\sigma_1^{(\mathrm{F})}}{\partial\varepsilon_2^{(\mathrm{Y})}}\mathbf{I}\otimes\mathbf{I} \quad \text{(S1.39)}$$

Note that we need $\partial\sigma_i^{(\mathrm{F})}/\partial\varepsilon_j^{(\mathrm{Y})}$ for all three cases which can be determined by taking the first derivative of $\sigma_i^{(\mathrm{F})}$ in (S1.30)

$$\frac{\partial\sigma_i^{(\mathrm{F})}}{\partial\varepsilon_i^{(\mathrm{Y})}} = \frac{\partial}{\partial\varepsilon_i^{(\mathrm{Y})}}\left(\frac{\partial f}{\partial\varepsilon_i^{(\mathrm{Y})}}\right) = \begin{cases} 0 & \varepsilon_i^{(\mathrm{Y})} < \epsilon_1 \\ \ell\,\dfrac{\left(\dfrac{\varepsilon_i^{(\mathrm{Y})} - \epsilon_1}{\epsilon_2 - \epsilon_1}\right)^t\left(\varepsilon_i^{(\mathrm{Y})} - \epsilon_1\right)}{t+1} & \epsilon_1 \le \varepsilon_i^{(\mathrm{Y})} < \epsilon_2 \\ \ell\left[\dfrac{\left(1 + \varepsilon_i^{(\mathrm{Y})} - \epsilon_2\right)^{s+1} - 1}{s+1} + \dfrac{\epsilon_2 - \epsilon_1}{t+1}\right] & \varepsilon_i^{(\mathrm{Y})} \ge \epsilon_2 \end{cases}$$

(S1.40)

With $\mathbf{C}^{(\mathrm{F})}$ at hand, the stiffness of the actin filament network $\mathbf{C}^{(\mathrm{A})}$ can be obtained from equation (S1.21).

### 2. Total cell stiffness

To determine the total stiffness of the cell, we first degrade the fourth-order tensors $\mathbf{C}^{(\mathrm{MT})}$ (S1.16), $\mathbf{C}^{(\rho)}$ (S1.19), and $\mathbf{C}^{(\mathrm{A})}$ (S1.21) to the second-order tensors $\mathbf{C}^{(\mathrm{MT})}$, $\mathbf{C}^{(\rho)}$, and $\mathbf{C}^{(\mathrm{A})}$. In Equation (1.20) we show how a fourth-order tensor (*e.g.*, $\mathrm{C}_{ijkl}^{(\rho)}$) is degraded to a second-order 6×6 matrix (*e.g.*, $\mathrm{C}_{ij}^{(\rho)}$).

Note that the microtubule network is connected to the myosin in parallel, and they are both connected to the actin network in series (Figure S17). Therefore, the total stiffness of the cell, $\mathbf{C}$, is obtained as follows

$$\mathbf{C} = \left( \left( \mathbf{C}^{(\mathrm{X})} \right)^{-1} + \left( \mathbf{C}^{(\mathrm{Y})} \right)^{-1} \right)^{-1} \tag{S1.41}$$

where

$$\mathbf{C}^{(\mathrm{X})} = \mathbf{C}^{(\rho)} + \mathbf{C}^{(\mathrm{MT})} \tag{S1.42}$$

and

$$\mathbf{C}^{(\mathrm{Y})} = \mathbf{C}^{(\mathrm{A})} = \mathbf{C}^{(\mathrm{I})} + \mathbf{C}^{(\mathrm{F})} \tag{S1.43}$$

### 3. Solving the set of nonlinear equations

In the previous sections, we presented the constitutive equations for the cytoskeletal model where the stress field $\sigma_{ij}$ is obtained from equation (S1.17) or (S1.25) and the stiffness field $\mathrm{C}_{ij}$ is determined from (S1.41). However, note that the stress and stiffness tensors $\sigma_{ij}$ and $\mathrm{C}_{ij}$ are functions of the unknown strain tensors $\varepsilon_{ij}^{(\mathrm{X})}$ and $\varepsilon_{ij}^{(\mathrm{Y})}$ (and not $\varepsilon_{ij}$). To determine $\varepsilon_{ij}^{(\mathrm{X})}$ and $\varepsilon_{ij}^{(\mathrm{Y})}$, we first define the following 12×1 vector

$$\begin{aligned}\boldsymbol{u} &= \left\{ \varepsilon_{11}^{(\mathrm{X})} \quad \varepsilon_{22}^{(\mathrm{X})} \quad \varepsilon_{33}^{(\mathrm{X})} \quad \varepsilon_{12}^{(\mathrm{X})} \quad \varepsilon_{13}^{(\mathrm{X})} \quad \varepsilon_{23}^{(\mathrm{X})} \quad \varepsilon_{11}^{(\mathrm{Y})} \quad \varepsilon_{22}^{(\mathrm{Y})} \quad \varepsilon_{33}^{(\mathrm{Y})} \quad \varepsilon_{12}^{(\mathrm{Y})} \quad \varepsilon_{13}^{(\mathrm{Y})} \quad \varepsilon_{23}^{(\mathrm{Y})} \right\}^{\mathrm{T}} \\ &= \{ u_1 \quad u_2 \quad \cdots \quad u_{12} \}^{\mathrm{T}}\end{aligned} \tag{S1.44}$$

which contains all 12 unknown variables in the strain tensors $\varepsilon_{ij}^{(\mathrm{X})}$ and $\varepsilon_{ij}^{(\mathrm{Y})}$. To determine the 12 unknowns, we need 12 equations. As the actin filament is connected to other elements in series (Figure S17), we use the following condition which gives us 6 equations

$$\boldsymbol{\sigma} = \boldsymbol{\sigma}^{(\mathrm{X})} = \boldsymbol{\sigma}^{(\mathrm{Y})} \tag{S1.45}$$

where the stress $\boldsymbol{\sigma}^{(\mathrm{X})}$ (equations (S1.17) and (S1.18))

$$\sigma_{ij}^{(\mathrm{X})} = \sigma_{ij} = \left( \mathrm{C}_{ijkl}^{(\rho)} + \mathrm{C}_{ijkl}^{(\mathrm{MT})} \right) \varepsilon_{kl}^{(\mathrm{X})} + \bar{\rho}_0 \delta_{ij} \tag{S1.46}$$

is directly transmitted to the actin filament network $\boldsymbol{\sigma}^{(\mathrm{Y})}$ (equation (S1.41))

$$\sigma_{ij}^{(\mathrm{Y})} = \sigma_{ij} = \sigma_{ij}^{(\mathrm{A})} = \sigma_{ij}^{(\mathrm{I})} + \sigma_{ij}^{(\mathrm{F})} \tag{S1.47}$$

We get the other 6 equations from the following condition

$$\boldsymbol{\varepsilon} = \boldsymbol{\varepsilon}^{(\mathrm{X})} + \boldsymbol{\varepsilon}^{(\mathrm{Y})} \tag{S1.48}$$

where $\boldsymbol{\varepsilon}^{(\mathrm{X})}$ is the strain of the cytoskeletal components that are in compression (*e.g.*, microtubule network), $\boldsymbol{\varepsilon}^{(\mathrm{Y})}$ is the strain of the cytoskeletal components that are in tension (*e.g.*, actin filaments), and $\boldsymbol{\varepsilon}$ is the total strain of the cell (Figure S17). Note that all stress and strain tensors $\sigma_{ij}^{(\mathrm{X})}$, $\sigma_{ij}^{(\mathrm{Y})}$, $\varepsilon_{ij}^{(\mathrm{X})}$, and $\varepsilon_{ij}^{(\mathrm{Y})}$ are symmetric. Therefore, the conditions in (S1.45) and (S1.48) can be defined by the following 12 equations in the 12×1 vector $\boldsymbol{f}$

$$\boldsymbol{f} = \{f_1 \quad f_2 \quad \cdots \quad f_{12}\}^{\mathrm{T}} \tag{S1.49a}$$

where

$$f_1 = \sigma_{11}^{(\mathrm{X})} - \sigma_{11}^{(\mathrm{Y})} \tag{S1.49b}$$

$$f_2 = \sigma_{22}^{(\mathrm{X})} - \sigma_{22}^{(\mathrm{Y})} \tag{S1.49c}$$

$$f_3 = \sigma_{33}^{(\mathrm{X})} - \sigma_{33}^{(\mathrm{Y})} \tag{S1.49d}$$

$$f_4 = \sigma_{12}^{(\mathrm{X})} - \sigma_{12}^{(\mathrm{Y})} \tag{S1.49e}$$

$$f_5 = \sigma_{13}^{(\mathrm{X})} - \sigma_{13}^{(\mathrm{Y})} \tag{S1.49f}$$

$$f_6 = \sigma_{23}^{(\mathrm{X})} - \sigma_{23}^{(\mathrm{Y})} \tag{S1.49g}$$

$$f_7 = \varepsilon_{11} - \varepsilon_{11}^{(\mathrm{X})} - \varepsilon_{11}^{(\mathrm{Y})} \tag{S1.49h}$$

$$f_8 = \varepsilon_{22} - \varepsilon_{22}^{(\mathrm{X})} - \varepsilon_{22}^{(\mathrm{Y})} \tag{S1.49i}$$

$$f_9 = \varepsilon_{33} - \varepsilon_{33}^{(\mathrm{X})} - \varepsilon_{33}^{(\mathrm{Y})} \tag{S1.49j}$$

$$f_{10} = \varepsilon_{12} - \varepsilon_{12}^{(\mathrm{X})} - \varepsilon_{12}^{(\mathrm{Y})} \tag{S1.49k}$$

$$f_{11} = \varepsilon_{13} - \varepsilon_{13}^{(\mathrm{X})} - \varepsilon_{13}^{(\mathrm{Y})} \tag{S1.49l}$$

$$f_{12} = \varepsilon_{23} - \varepsilon_{23}^{(\mathrm{X})} - \varepsilon_{23}^{(\mathrm{Y})} \tag{S1.49m}$$

We then determine the 12×12 Jacobian matrix $\mathbf{J}$

$$\mathbf{J}=\begin{bmatrix}\partial f_1/\partial u_1 & \partial f_1/\partial u_2 & \dots & \partial f_1/\partial u_{12}\\ \partial f_2/\partial u_1 & \partial f_2/\partial u_2 & \dots & \partial f_2/\partial u_{12}\\ \vdots & \vdots & & \vdots\\ \partial f_{12}/\partial u_1 & \partial f_{12}/\partial u_2 & \dots & \partial f_{12}/\partial u_{12}\end{bmatrix}=\begin{bmatrix}\mathbf{C}^{(\mathrm{X})} & -\mathbf{C}^{(\mathrm{Y})}\\ -\mathbf{I} & -\mathbf{I}\end{bmatrix} \tag{S1.50}$$

using equations (S1.44) and (S1.49) for $u_i$ and $f_i$, respectively. Finally, we use the Newton-Raphson method

$$\boldsymbol{u}_{i+1}=\boldsymbol{u}_i-\mathbf{J}^{-1}\boldsymbol{f}(\boldsymbol{u}_i) \tag{S1.51}$$

to determine the unknown vector $\boldsymbol{u}$, where the vectors $\boldsymbol{u}_i$ and $\boldsymbol{u}_{i+1}$ are respectively the solutions for $i$ and $i+1$ iterations. To find the solutions of the nonlinear equations, we use the following initial guess $\boldsymbol{u}_0$ (or any other initial guess)

$$\boldsymbol{u}_0=\{0 \quad 0 \quad \dots \quad 0\}^{\mathrm{T}}_{1\times 12} \tag{S1.52}$$

and the convergence criterion

$$|\boldsymbol{f}|=\sqrt{(f_1)^2+(f_2)^2+\cdots+(f_{12})^2}<\epsilon_{\mathrm{Tol}} \tag{S1.53}$$

where $|\boldsymbol{f}|$ is the magnitude of the vector $\boldsymbol{f}$, and $\epsilon_{\mathrm{Tol}}$ is the convergence threshold. Using $\epsilon_{\mathrm{Tol}} =$ $10^{-8}$ in our simulations, we stop the iterations in (S1.51) when $|\boldsymbol{f}|$ is less than $\epsilon_{\mathrm{Tol}}$. With the strain tensors $\varepsilon_{ij}^{(\mathrm{X})}$ and $\varepsilon_{ij}^{(\mathrm{Y})}$ determined from (S1.51), we can calculate the stress tensor $\sigma_{ij}$ (from (S1.17) or (S1.25)) and the stiffness tensor $\mathrm{C}_{ij}$ (from (S1.41)).

### 4. Actomyosin contractility increases with anisotropy in tension

Combining equation (S1.17) with (S1.18), one can derive the following expression for the average of contractility, $\frac{1}{3}\rho_{kk}=(\rho_{11}+\rho_{22}+\rho_{33})/3$

$$\frac{\rho_{kk}}{3}=\left(\frac{3K^{(\mathrm{MT})}\alpha_{\mathrm{v}}-1}{3K^{(\mathrm{MT})}\beta_{\mathrm{v}}-1}\right)\frac{\sigma_{kk}}{3}+\left(\frac{3K^{(\mathrm{MT})}\beta_{\mathrm{v}}}{3K^{(\mathrm{MT})}\beta_{\mathrm{v}}-1}\right)\rho_{0}, \tag{S1.54}$$

which shows that the average of contractility increases with the average of stress, $\frac{1}{3}\sigma_{kk}=(\sigma_{11}+\sigma_{22}+\sigma_{33})/3$. We have previously shown that the feedback mechanism between $\rho_{kk}$ and $\sigma_{kk}$ in (S1.54) predicts that contractility, cytoskeleton tension, and traction forces increase with substrate stiffness and substrate area, which are consistent with experimental observations (*55*).

As discussed in the main text, the model also accounts for the effect of tension anisotropy where phosphorylation of myosin increases with anisotropy in the components of the stress tensor $\sigma_{ij}$. The increase in actomyosin contractility in response to tension anisotropy is introduced to the model with the term $\alpha_{\mathrm{a}}\sigma_{\mathrm{a}}$

$$\frac{\rho_{kk}}{3}=\left(\frac{3K^{(\mathrm{MT})}\alpha_{\mathrm{v}}-1}{3K^{(\mathrm{MT})}\beta_{\mathrm{v}}-1}\right)\frac{\sigma_{kk}}{3}+\alpha_{\mathrm{a}}\sigma_{\mathrm{a}}+\left(\frac{3K^{(\mathrm{MT})}\beta_{\mathrm{v}}}{3K^{(\mathrm{MT})}\beta_{\mathrm{v}}-1}\right)\rho_{0} \tag{S1.55}$$

where $\sigma_1 > \sigma_2 > \sigma_3$ are the principal stress values (eigenvalues) of the stress tensor $\sigma_{ij}$ with $\sigma_1 > \sigma_2 > 0$, $\sigma_\mathrm{a} = \tanh\left(\frac{1}{2}\left(\frac{\sigma_1}{\sigma_2} - 1\right)\right)\sigma_1$ represents the tension anisotropy, $\alpha_\mathrm{a}$ is the anisotropic chemo-mechanical feedback parameter which regulates the increase in myosin phosphorylation with tension anisotropy. The additional term $\alpha_\mathrm{a}\sigma_\mathrm{a}$ is implemented into the finite element framework using a piecewise linear approximation where $\rho_0$ in equation (S1.13) is simply replaced with $\rho_0 + \left(\frac{3K^{(\mathrm{MT})}\beta_\mathrm{v} - 1}{3K^{(\mathrm{MT})}\beta_\mathrm{v}}\right)\alpha_\mathrm{a}\sigma_\mathrm{a}$ in each step of the simulation.

**Supplementary Figures**

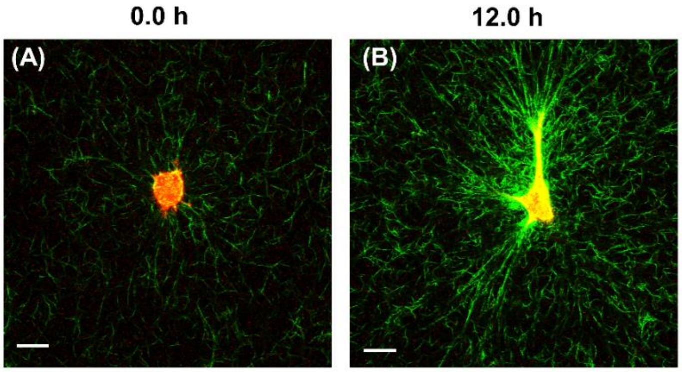


**Figure S1. Fibroblast transformation within collagen fibrous microenvironments.** To track the transformation of cells, we used live-cell microscopy where fluorescence confocal imaging of cells labeled with Cell Tracker Orange and confocal reflectance imaging of collagen in 3D polymerized collagen matrices were simultaneously done on a microscope using a water immersion objective (see Methods). (A) Fibroblasts were initially round, small, and inactive, (B) and they became polarized, spread, and contractile several hours after being cultured within their matrices. Scale bar: 20 µm

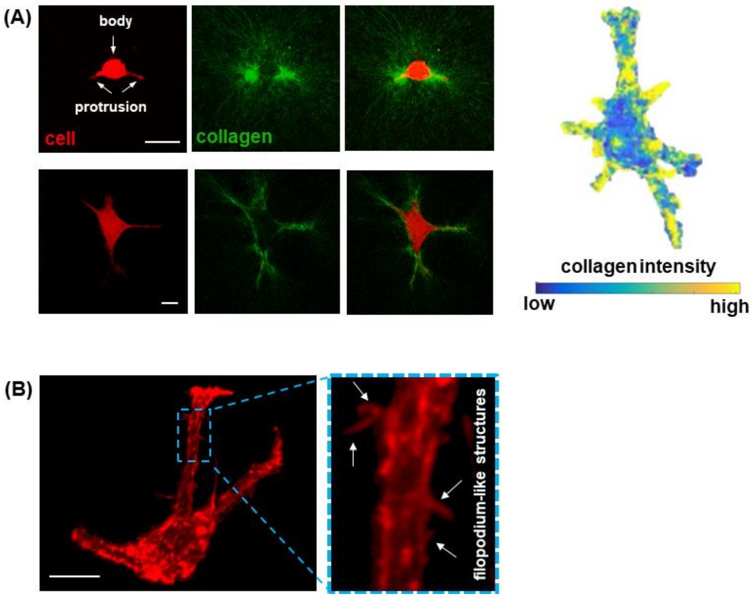


**Figure S2. Fibroblasts used protrusions (and not the cell body) as the primary means to accumulate collagen.** (A) In the early stages of cell-matrix interaction, compared to the body of the cell, protrusions aligned and accumulated substantially more collagen fibers, indicating that cells used their protrusions to generate tension and transform themselves from their initial inactive state to an active contractile state. (B) These protrusions had small actin-based filopodium-like structures that may facilitate the adhesion between protrusions and collagen fibers (*38*). Scale bar: 20 μm

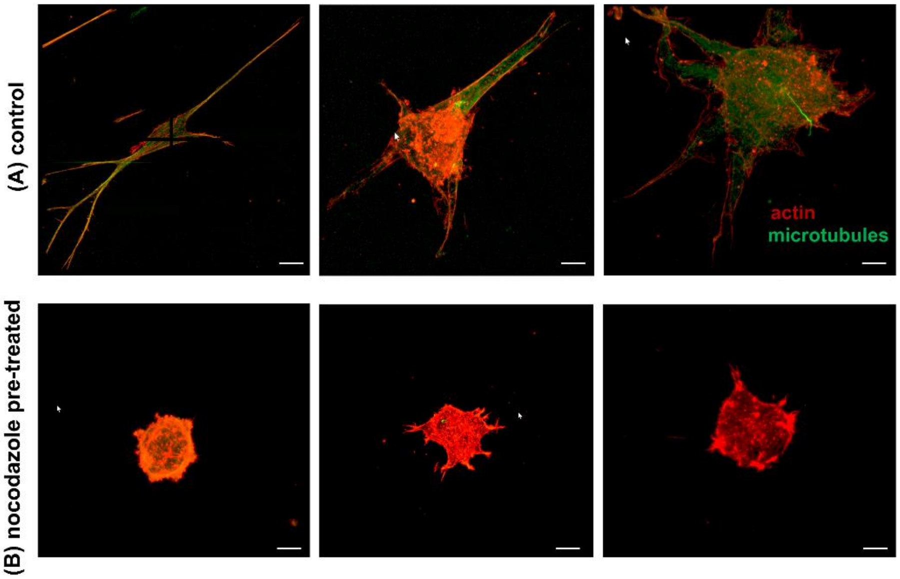


**Figure S3. Microtubules form the core of cellular protrusions.** Unlike control cells (A), fibroblasts with pre-disrupted microtubules were not able to form protrusions at any stage of cell-matrix interactions and they remained round (B). Scale bar: 10 µm

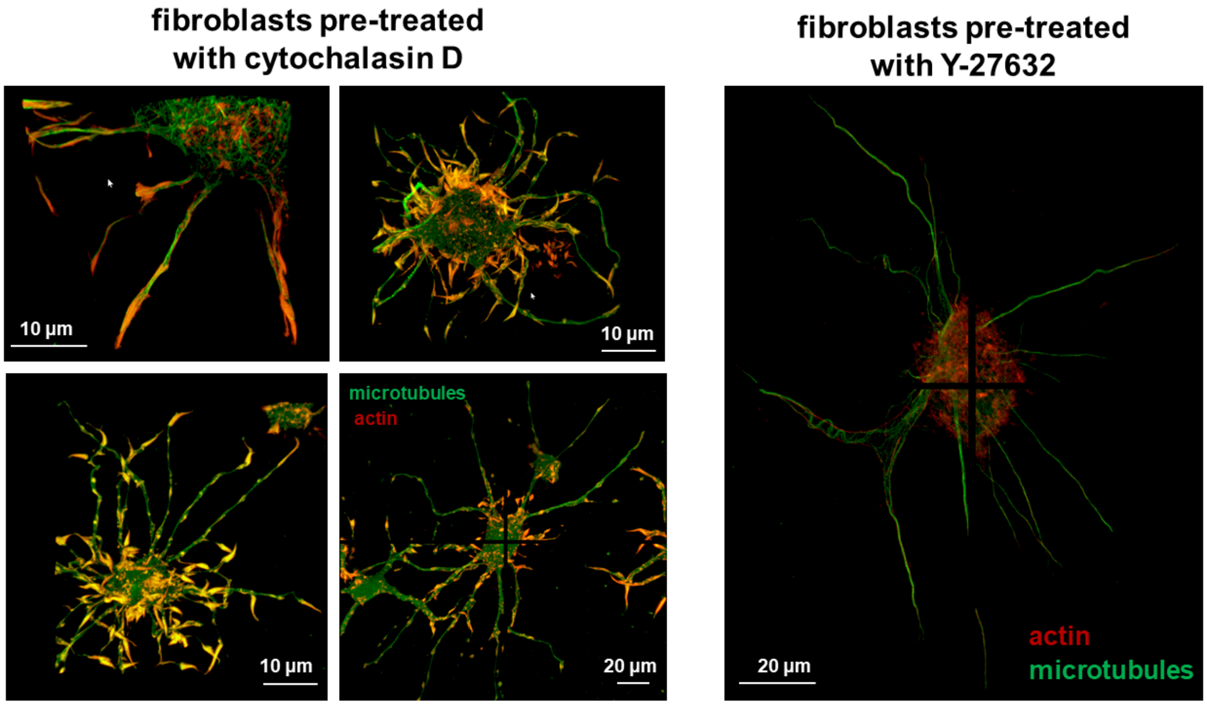


**Figure S4. Pre-treatment of fibroblasts with inhibitors of actomyosin contractility further showed the importance of microtubules in the formation of cellular protrusions.** Pre-treatment of cells with actomyosin inhibitors (cytochalasin D and Y-27632) grew cellular protrusions in length and number where microtubules formed the core of these protrusions (*38*).

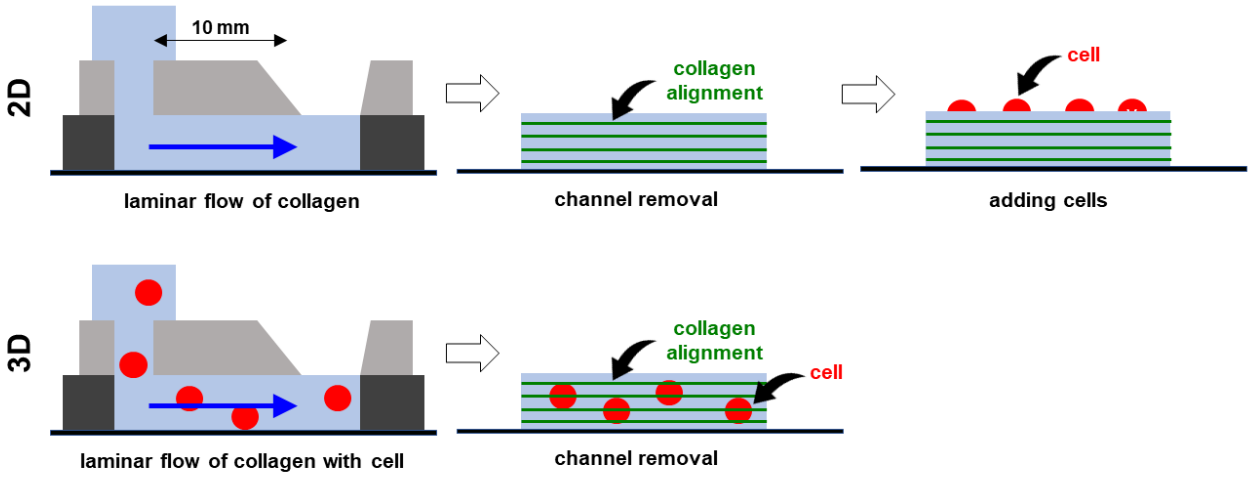


**Figure S5. Generation of two-dimensional and three-dimensional tissues with aligned collagen fibers.** Schematics of the micro fluid channel.

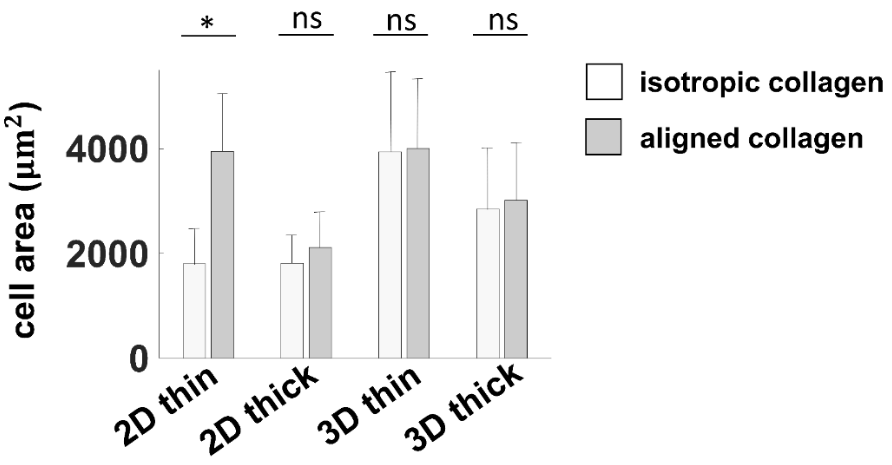


**Figure S6. The spreading of fibroblasts in 3D tissues is independent of the matrix fiber organization.** Fibroblasts spread well atop and within both isotropic and aligned ECMs. Except on thin, 2D substrata, cell spreading area was independent of matrix fiber organization as fibroblasts showed almost the same spreading area in isotropic and aligned ECMs. Error bars represent standard deviations ($n$ = 39-104).

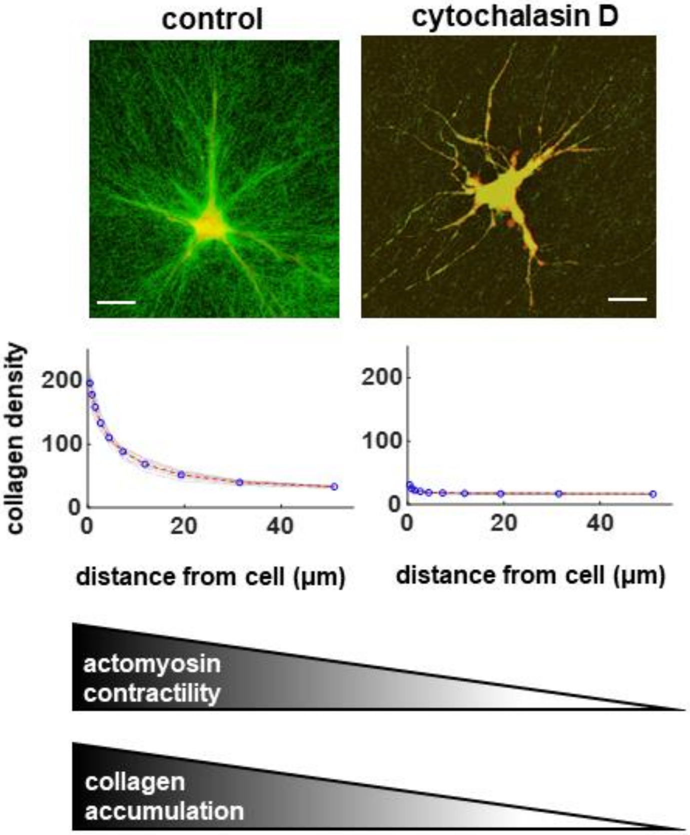


**Figure S7. Collagen accumulation and alignment required actomyosin contractility.** Cellular protrusions grew in length and number upon pre-treatment of cells with actomyosin inhibitors. However, these protrusions did not align and accumulate collagen fibers. Scale bar: 20 μm

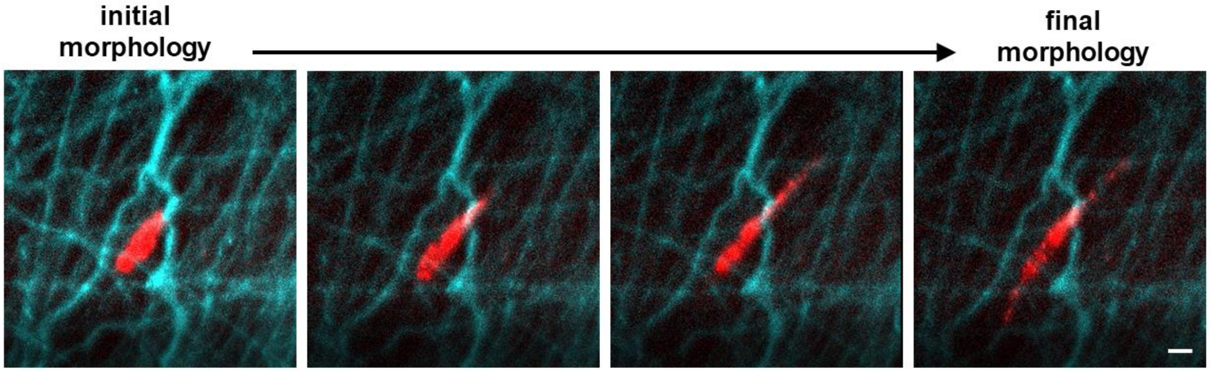


**Figure S8. Polarization of fibroblasts in the direction of pre-aligned fibers.** Transformation of fibroblasts within three-dimensional electrospun synthetic fibrous hydrogels. The white arrow shows the predominant direction of pre-aligned synthetic hydrogel fibers. Scale bar: 20 µm

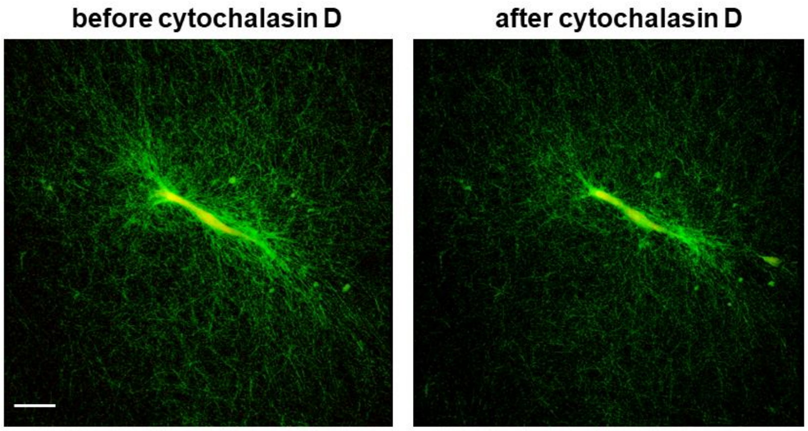


**Figure S9. Cell contraction induces both elastic and plastic deformations in the surrounding collagen fibrous matrix.** Fibroblasts were post-treated with cytochalasin D to decrease cell actomyosin contractility. Upon cytochalasin D, collagen fibers were relaxed and collagen accumulation significantly decreased which indicates the elastic nature of cell-induced matrix deformation. However, some degrees of collagen alignment can be still observed which may indicate the existence of cell-induced plastic deformation in the matrix. Scale bar: 20 µm

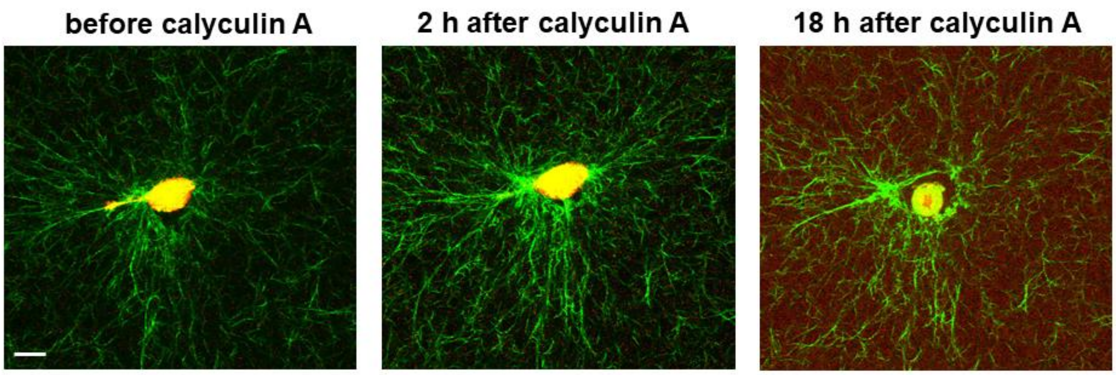


**Figure S10. Cell-induced plastic deformations in collagen fibrous matrices.** To further investigate the existence of cell-induced plastic deformation in collagen matrices, fibroblasts were post-treated with calycullin A to increase cell actomyosin contractility. The sudden increase in cell contractility led to significant cell shrinkage which caused detachment of the cell from the surrounding matrix. Even after detachment of the cell from the matrix, some collagen fibers were remained aligned indicating that, in addition to elastic deformation, cell contraction can also generate plastic deformation in the matrix. Scale bar: 20 μm

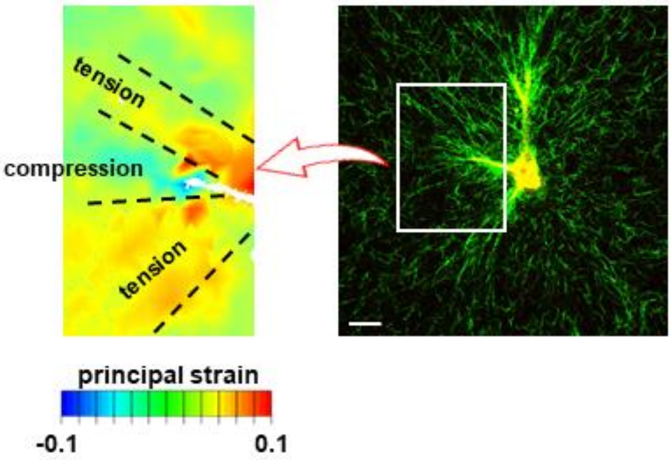


**Figure S11. Fibroblast protrusions grew from the tip while the protrusion side maintained the tension at the cell-matrix interface.** We used a three-dimensional strain mapping technique to determine the cell-induced strain field around cell protrusions. As expected, with the growth of a protrusion, we found compressive strains in a cone ahead of the protrusion. However, our results showed that, while the protrusion was growing from the tip, it pulled on the matrix by the lateral sides to maintain the tension at the cell-matrix interface. We used 20 min time increment for the strain field measurement. Scale bar: 20 µm

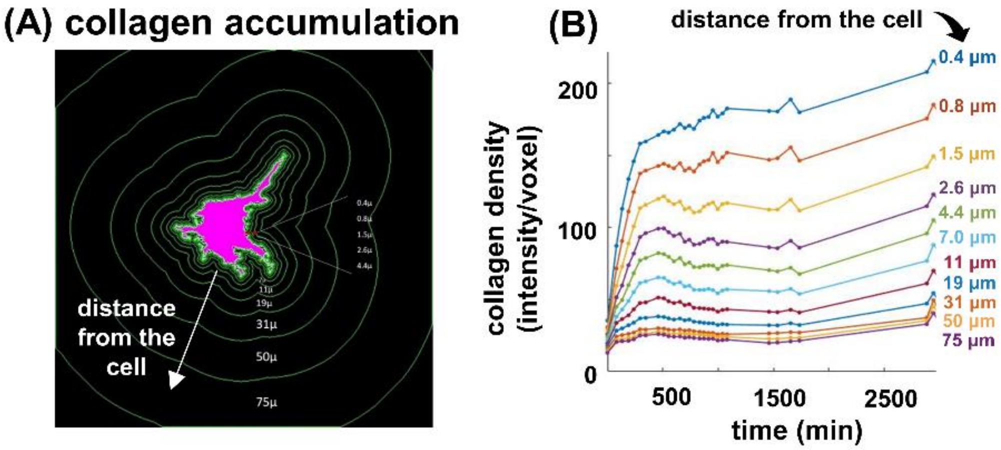


**Figure S12. Collagen accumulation reached a plateau over time.** To measure collagen accumulation around a cell, we performed confocal fluorescence and reflectance imaging in three-dimensional volumetric regions surrounding each cell at various time points. We then quantified the degree of collagen accumulation using a custom MATLAB script that averaged the confocal reflectance pixel intensity within volumetric shells surrounding each cell (A). The highest level of collagen accumulation was observed near the cell (blue line in (B) with 0.4 μm distance from the cell membrane). Also, our results showed that collagen accumulation in all surrounding regions reached a plateau over a time scale of hundreds of minutes. The collagen density in different regions shown in (A) was quantified in (B) where the distance of each region from the cell membrane was presented next to its curve.

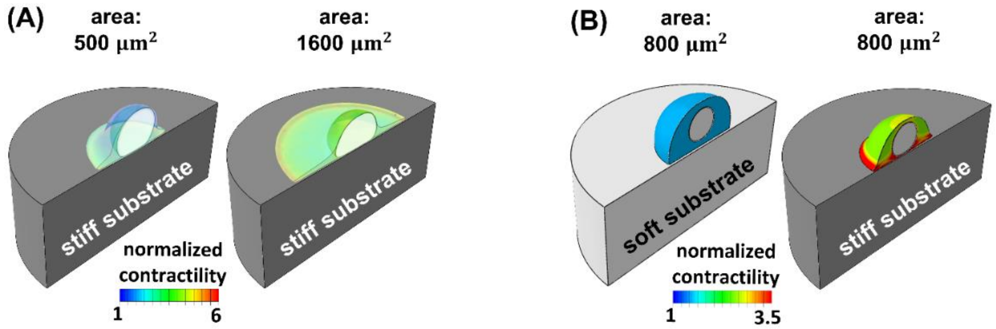


**Figure S13. Cells sense and respond to mechanical signals from the extracellular matrix.** Cell contractility (denoted by the average density of phosphorylated myosin motors) increases with increasing external resistance from the matrix which can be caused by increasing cell-substrate contact area (A) or increasing substrate stiffness (B). As we have previously shown (*55*), the increase in cell contractility is in agreement with experimental observations where cells cultured on micropatterned substrates with larger areas (or stiffer matrices) exhibited higher levels of phosphorylated myosin and generated higher contractile forces compared with cells cultured on micropatterned substrates with smaller areas (or softer matrices).

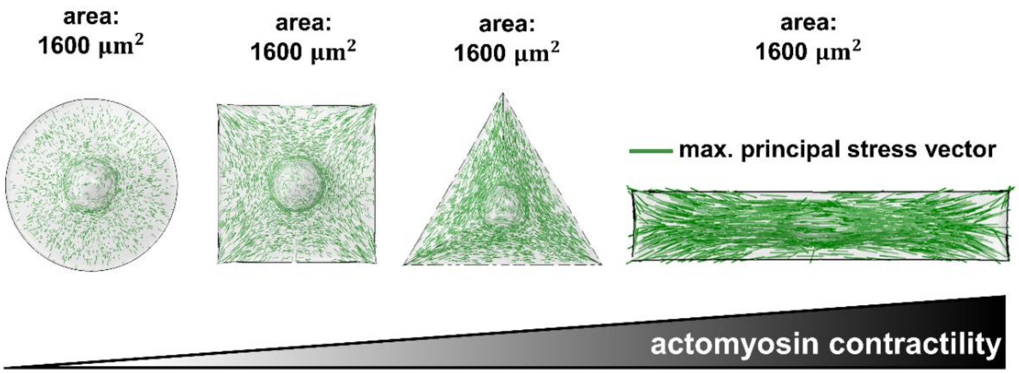


**Figure S14. Anisotropy of the cytoskeletal stress field promotes cell actomyosin contractility.** Simulations of fibroblast cells cultured on rigid micropatterned substrates with a surface area of 1600 $\mu m^2$ showed that cell actomyosin contractility increased with the degree of substrate polarization. In addition to the magnitude of tension (regulated by substrate area and substrate stiffness as shown in Figure S13), the polarity of the tensile field (tension anisotropy) also impacted the level of cell contractility. Cells on substrates with higher degrees of polarization experienced higher degrees of tension anisotropy leading to higher actomyosin contractility and generation of higher contractile forces, which is consistent with experimental observations as we have reported before (*55*).

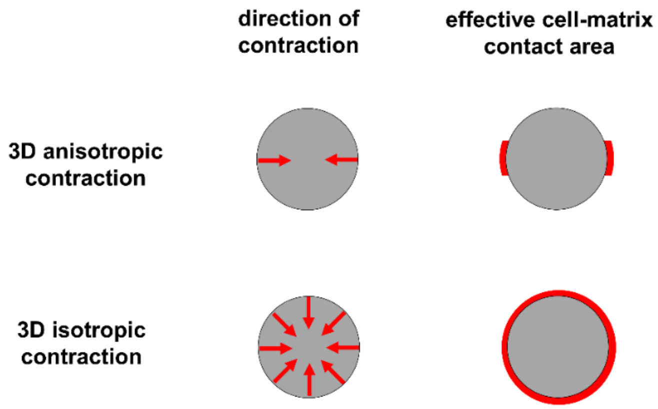


**Figure S15. The effective cell-matrix contact area in polarized contraction is significantly lower than isotropic contraction.** Compared with isotropic contraction, when a cell generates polarized contraction, a lower surface area of the cell is exposed to the resistance from the surrounding matrix.

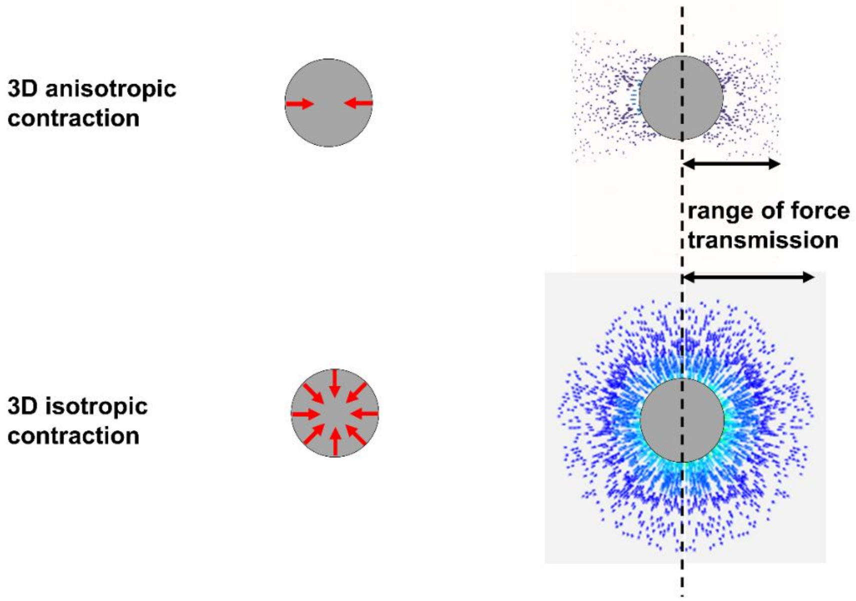


**Figure S16. The effect of tension anisotropy term on the modeling results.** Cell models that do not account for the effect of tension anisotropy (e.g., pre-stressed/pre-strained cell models and thermoelastic-based classical cell models) predict that cells experience higher cytoskeletal tension, generate higher contractile forces and collagen alignment if they contract isotropically, which are opposite to our model predictions in Fig. 5.

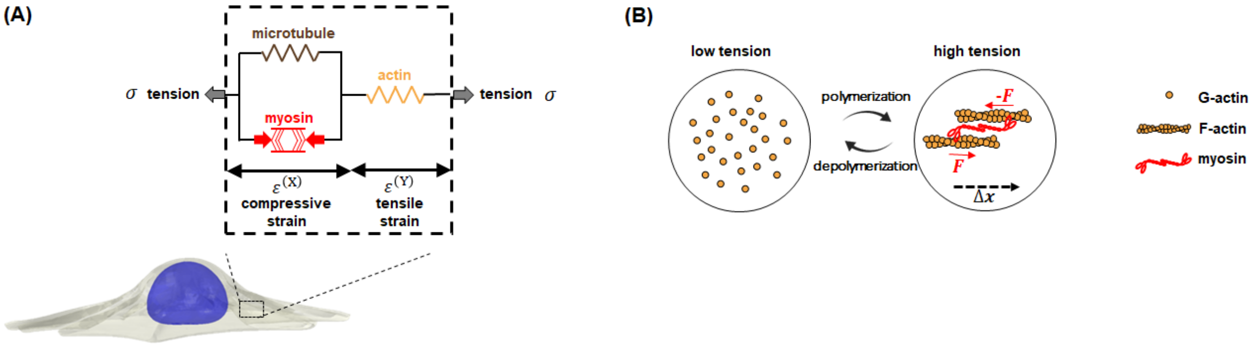


**Figure S17. Components of the cytoskeletal model.** (A) The cytoskeleton is composed of (i) the myosin motors, (ii) the microtubule network, and (iii) the actin filament network. (B) In response to the tension $\sigma$ (generated either by the intrinsic cell actomyosin contractility or external tensile forces) the cell contractility $\rho$ and the stiffness of the actin element $C^{(A)}$ increase with tension, representing recruitment of myosin motors and polymerization of actin filaments, respectively.

### Supplementary Movie

**Movie S1. Cell alignment along the direction of pre-aligned ECM fibers.** Growth of cell protrusions in the direction of synthetic ECM fibers.
